# Benzoxaboroles are structurally unique binders of eukaryotic translation initiation factor 4E

**DOI:** 10.64898/2026.02.24.707563

**Authors:** Joshua B. Combs, D. Matthew Peacock, Gregory B. Craven, Sungwon Jung, Ying Chen, Sang M. Le, Jack Taunton, Kevan M. Shokat

## Abstract

Benzoxaboroles offer unusual reactivity and protein recognition for the development of small molecule drugs. Despite this potential, they are uncommon in drug discovery or in fragment screening libraries. We synthesized a series of structurally related benzoxaboroles containing a diazirine/alkyne tag to enable in-cell photoaffinity labeling experiments. A subset of this library was found to have high selectivity for eukaryotic translation initiation factor 4E (eIF4E). The benzoxaborole–eIF4E interaction was found to be stereoselective in nature and competitive with the 7-methylguanosine cap of mRNA. Site of labeling experiments revealed that the benzoxaborole fragment interacts with the cap binding pocket of eIF4E. *In silico* modeling of the modified protein suggests that H-bonding interactions between the main chain of Trp102 and the side chain of Asn155 to the amide carbonyl and anionic boronate of the benzoxaborole, respectively, drive affinity for this challenging to drug pocket.

---

Benzoxaborole inhibitors target an array of proteins through diverse modes of interaction. As a neutral species they can interact with the hinge motif of kinases^1^, and as the anionic boronate they can coordinate Lewis acidic metal atoms^2, 3^, covalently bind to serine and threonine residues^4^, and perhaps most uniquely the 2’,3’ *cis*-diol ribose sugar at the 3’ end of RNA^5, 6^ (Figure 1a).^7-12^ The two FDA-approved benzoxaboroles, tavaborole and crisaborole, are approximately half the molecular weight of FDA-approved compounds from a similar period, suggesting they possess high ligand efficiency and significant potential for unique protein recognition properties.^2, 5, 13^

**Figure 1.**
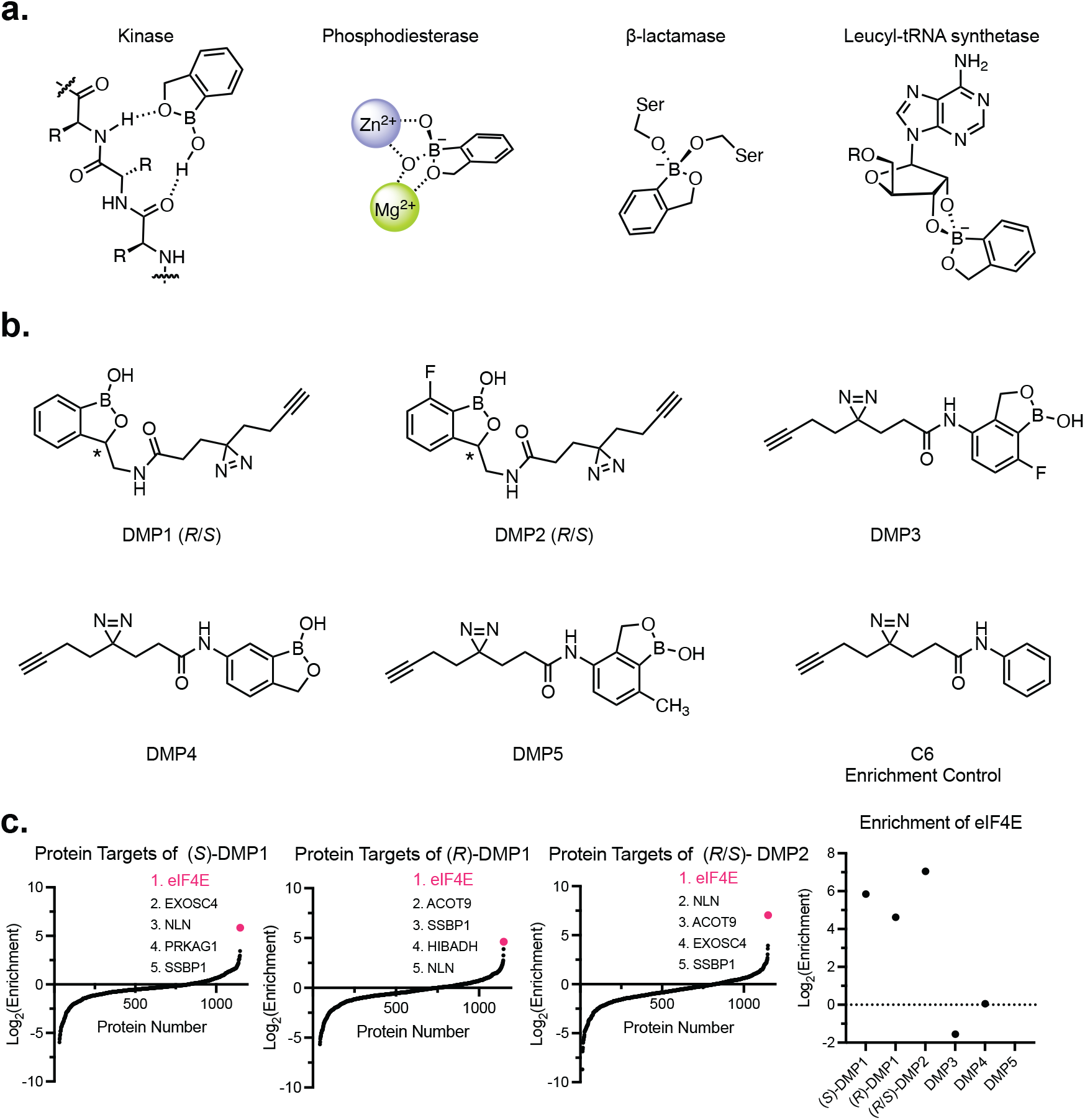
(a) Precedented modes of protein binding for benzoxaboroles. (b) Benzoxaborole fragment series and enrichment control. (c) log_2_[Enrichment] vs protein number for fragments with high enrichment of eIF4E. Protein number corresponds to the target’s position in a list of identified proteins. No value is reported for **DMP5** because no TMT signal was observed.

Due to their interesting physical chemical properties and ability to bind proteins with unique mechanisms, we synthesized a series of benzoxaborole compounds that contain a photoaffinity labeling (PAL) probe comprising a photoreactive diazirine and ‘click’-chemistry enabling alkyne.^14^ The PAL probe was attached either at the 3-position of the oxaborole ring (**DMP1-2**) or at positions on the aromatic ring (**DMP3-5**), to determine if installation at different sites would affect protein enrichment (Figure 1b). HEK293T cells were treated with each compound, photoirradiated (365 nm), and lysed. A biotin ‘click’-reaction on the lysates allowed for enrichment of compound labeled proteins with streptavidin affinity purification. Digestion of the captured proteins enabled identification of peptide fragments through quantitative MS-based proteomics. Enrichment was calculated by normalizing the total TMT reporter intensities for each probe, then dividing the TMT intensity for a compound by the TMT intensity for the control probe **C6**.^15, 16^

**(*S*)-** and **(*R*)-DMP1** and racemic **DMP2** showed high levels of enrichment for eIF4E (Figure 1c). Even though both enantiomers of **DMP1** were found to label eIF4E, **(*S*)-DMP1** had a >2x fold increased enrichment value for eIF4E versus **(*R*)-DMP1**. Substitution of a fluorine atom *ortho*- to the boron-carbon bond (**DMP2**) has been shown to reduce the p*K*_a_ of the benzoxaborole and also enhanced eIF4E enrichment.^17^ Attachment of the PAL warhead through the aryl ring resulted in diminished enrichment values (log_2_[Enrichment]<0) (**DMP3** and **DMP4**), or no detectable eIF4E labeling (**DMP5**). These results are consistent with prior reports on enantiomeric probe pairs and suggest that **DMP1** and **DMP2**’s target enrichment of eIF4E is not a universal characteristic of benzoxaboroles.^18, 19^

Recognition of the 5’-7-methylguanosine (m7G) cap of eukaryotic mRNA by the eIF4E component of the eukaryotic initiation factor 4F (eIF4F) protein complex is thought to be the rate limiting step of eukaryotic translation.^20, 21^ In 1990, Sonenberg and co-workers reported that overexpression of eIF4E in cells led to malignant transformation, and subsequent studies have shown that excess eIF4E can be important for the growth of tumors in transgenic mice.^22, 23^ While there are no FDA approved inhibitors of eIF4E, several classes of translation inhibitors have been developed that act both directly and indirectly on it.^24^ Cap competitive eIF4E inhibitors that mimic the structure of m7G have been reported as potential efficacious agents, but cell permeability has been a common problem.^25-29^ Second generation cap analogues have begun to address cell penetration by utilizing bis-POM phosphonate esters as prodrugs and display increased cell activity.^30^ Diazirine probes have also been found to have modest eIF4E enrichment and have been mapped to a peptide fragment adjacent to the m7G binding pocket (Supplementary Figure 1).^31^ Rapamycin has been shown to impact activation of eIF4E’s inhibitory partner protein, eukaryotic translation initiation factor 4E-binding protein. This antagonism of eIF4E is thought to play a role in Rapamycin’s suppression of tumor growth.^32, 33^

In-gel fluorescence and western blot experiments were used to validate eIF4E as a target of **(*S*)-DMP1** and **DMP2**. A copper-catalyzed ‘click’ reaction with TAMRA Azide Plus on compound labeled cell lysates allowed in-gel visualization (550 nm) of labeled proteins. Intense TAMRA bands were observed at the appropriate size for eIF4E when cells were treated with 100 µM **(*S*)-DMP1** or **(*S*)-DMP2** (Figure 2a, Supplementary Figure 2). Independently blotting for eIF4E with commercial rabbit and mouse antibodies displayed an unexpected phenomenon. Immunoblots with the rabbit-derived antibody contained higher intensity bands from samples labelled with **(*S*)-DMP1** or **(*S*)-DMP2**, compared to the DMSO control. However, lower intensity bands were observed from the same samples when probed with the mouse derived eIF4E antibody. A labeling dose response was used to further interrogate the effect **(*S*)-DMP2** had on antibody recognition. Cells treated with a range of concentrations (0–100 µM) of **(*S*)-DMP2** showed a dose-dependent increase in labeling by in-gel fluorescence, and the same rabbit vs mouse antibody recognition trend was also observed (Figure 2b). These results are consistent with labeling of eIF4E by **(*S*)-DMP2** and suggest **(*S*)-DMP2**’s site of labeling disrupts the epitope recognized by the mouse antibody.^34^

**Figure 2.**
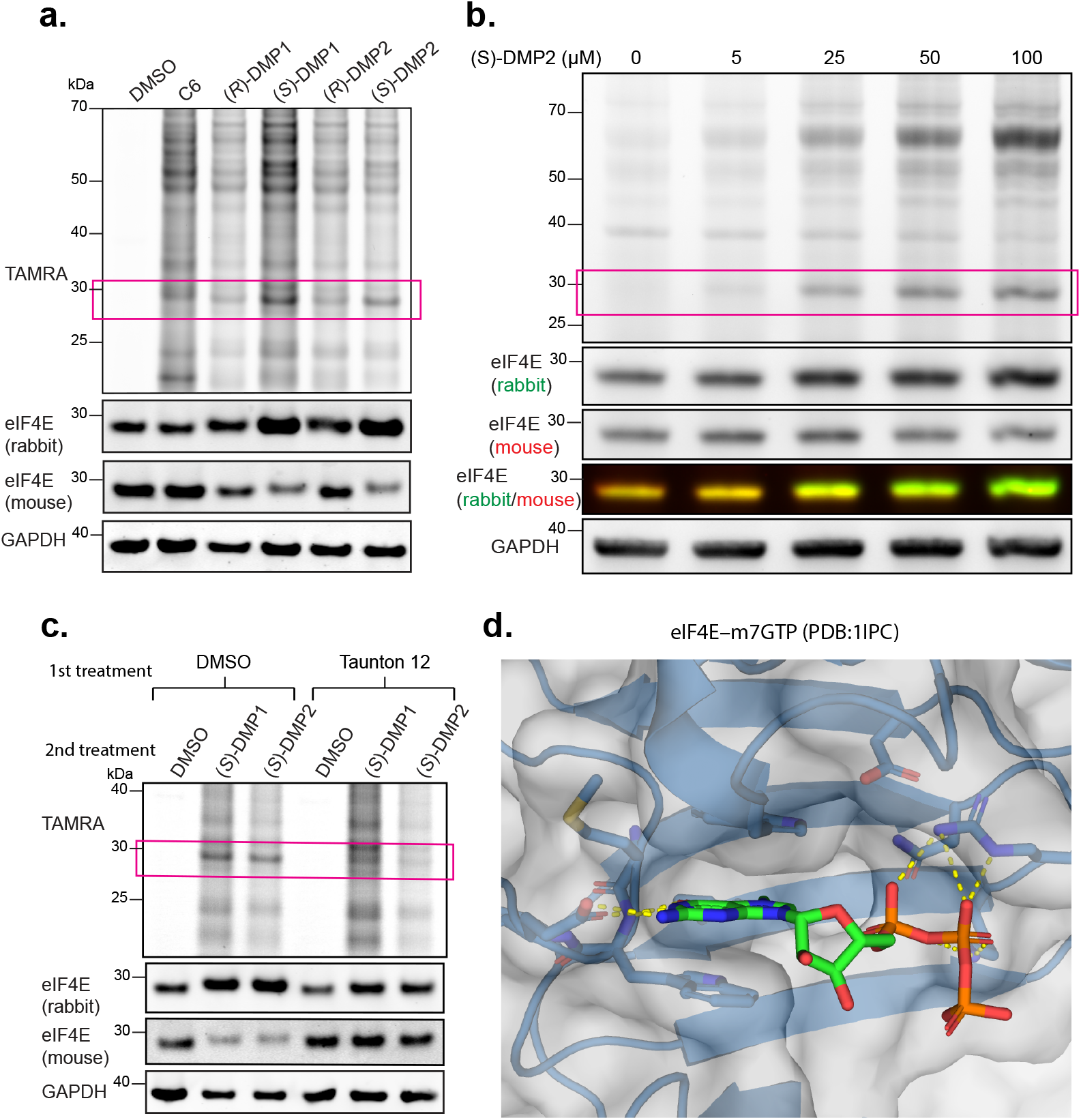
(a) In-gel fluorescence and western blot from HEK293Ts treated with 100 µM compound. The boxed bands are at the gel shift for labeled eIF4E. (Triplicate) (b) A dose curve of (*S*)-DMP2. (Duplicate) (c) Labeling competition with covalent eIF4E inhibitor **Taunton 12**. (Triplicate) (d) Co-crystal structure of m7GTP bound to eIF4E. (PDB: 1IPC).^35^

A competitive labeling experiment with the covalent eIF4E inhibitor **Taunton 12** was used to determine if photo-crosslinking of **(*S*)-DMP1** and **(*S*)-DMP2** to eIF4E would be prevented by occupation of the 5’-cap binding site.^25^ **Taunton 12** contains a sulfonyl fluoride moiety that reacts with K162 and blocks binding of m7G derivatives. HEK293T cells were incubated with DMSO or 10 µM **Taunton 12** before treatment with DMSO, **(*S*)-DMP1**, or **(*S*)-DMP2**. Compared to the DMSO control, the in-gel fluorescence for cells treated with **Taunton 12** no longer had a band at the expected size for eIF4E. The anti-eIF4E western blot also showed that recognition by the mouse antibody was restored in the samples pretreated with **Taunton 12** (Figure 2c). This result suggests that while **Taunton 12** and **DMP1/2** bind at the same pocket, the site(s) of covalent modification by **DMP1/2** following photoirradiation are distinct from the K162 modified by **Taunton 12**.

Based on available crystal structures of eIF4E, recognition of the 7-methylguanosine cap of mRNA is driven by the 5’-methyl guanine nucleoside and triphosphate moiety.^35^ The ribose diol is solvent exposed, posing two possible mechanisms based on established benzoxaborole modes of binding (Figure 1a). The benzoxaboroles could be directly competitive with the nucleoside ligand as reported for the ATP competitive kinase inhibitor **AN3484**, or they could covalently engage the diol of the 7-methylguanisine ligand as reported for the leucyl-tRNA synthe-tase inhibitor tavaborole (Figure 2d).^1, 6^

To differentiate between these two mechanisms of protein recognition, **(*S*)-DMP2** (1–50 µM) was incubated with recombinantly expressed eIF4E (28–217) and then photoirradiated. The TAMRA-’click’ procedure was used as a read out for the relative amount of ligated protein. Compared to incubation of eIF4E with only **(*S*)-DMP2**, addition of the cap mimic m^7^G(5′)ppp(5′)G (**m7GTPG**) or **Taunton 12** led to decreased amounts of labeled protein (Figure 3a). To control for possible sequestration of the benzoxaborole by the cap mimic leading to decreased labeling, a non-cap competitive G(5′)ppp(5′)G (**GTPG**) control was used.^36^ **GTPG** had a minimal impact on **(*S*)-DMP2** labeled eIF4E indicating that the reduced labeling in the presence of **m7GTPG** and **Taunton 12** was due to occupation of the cap binding motif of eIF4E. A control for general diazirine reactivity can be found in Supplementary Figure 3.

**Figure 3.**
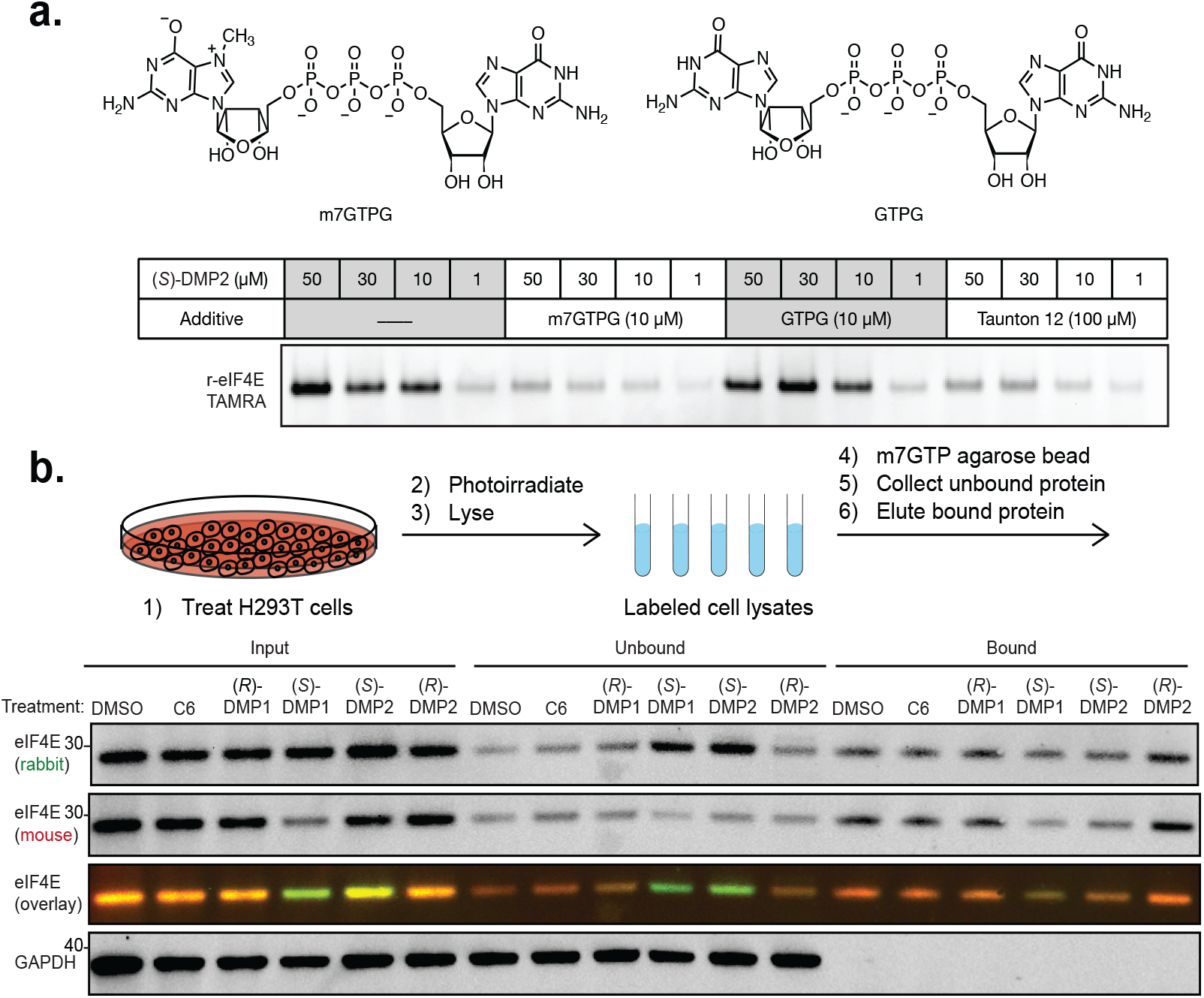
(a) Labeled recombinant eIF4E (28–217) visualized by in-gel fluorescence. Labeling was performed with no additive, m7GTPG, GTPG, and Taunton 12. (Duplicate) (b) Active eIF4E was chemoprecipitated from cell lysate utilizing an agarose bead modified with m7GTP. A western blot with rabbit and mouse anti-eIF4E antibodies was used to probe for eIF4E in the input, the unbound fraction, and the bound fraction of lysate. (Triplicate, quantified in Supplementary Figure 4)

While it was possible to establish that binding of **m7GTPG** was competitive with protein labeling by the benzoxaborole probes, we also wanted to establish whether in-cell engagement of eIF4E perturbed cap binding. The same treatment and photoirradiation protocol that was used in the chemoproteomics experiments was used to generate compound labeled cell lysates. A commercial agarose bead functionalized with m7GTP was used to affinity purify cap binding proteins from cell lysates. After collecting the unbound lysate fraction, the beads were washed, and bound proteins eluted (Figure 3b).^25^ The eIF4E from cells treated with **(*S*)-DMP1** and **(*S*)-DMP2** displayed decreased binding to the cap-functionalized beads based on the increase in eIF4E found in the unbound protein sample and decrease in the bound sample (Supplementary Figure 4). The ratio of eIF4E recognized by the rabbit versus mouse antibody also supports that there is an enrichment of benzoxaborole-labeled protein in the unbound fractions.

To gain deeper insights into how **(*S*)-DMP2** binds eIF4E, we mapped the sites of diazirine insertion into eIF4E.^31, 37, 38^ **(*S*)-DMP2** labeled protein was digested with trypsin and subjected to mass spectrometry-based proteomics identifying two **(*S*)-DMP2** modified peptides. The modified peptides were identical in sequence (D96–K106) and matched the peptide identified in previous diazirine containing ligands with modest eIF4E enrichment. Met101 was modified in one instance while Glu103 was modified in the other (Figure 4a/b).

**Figure 4.**
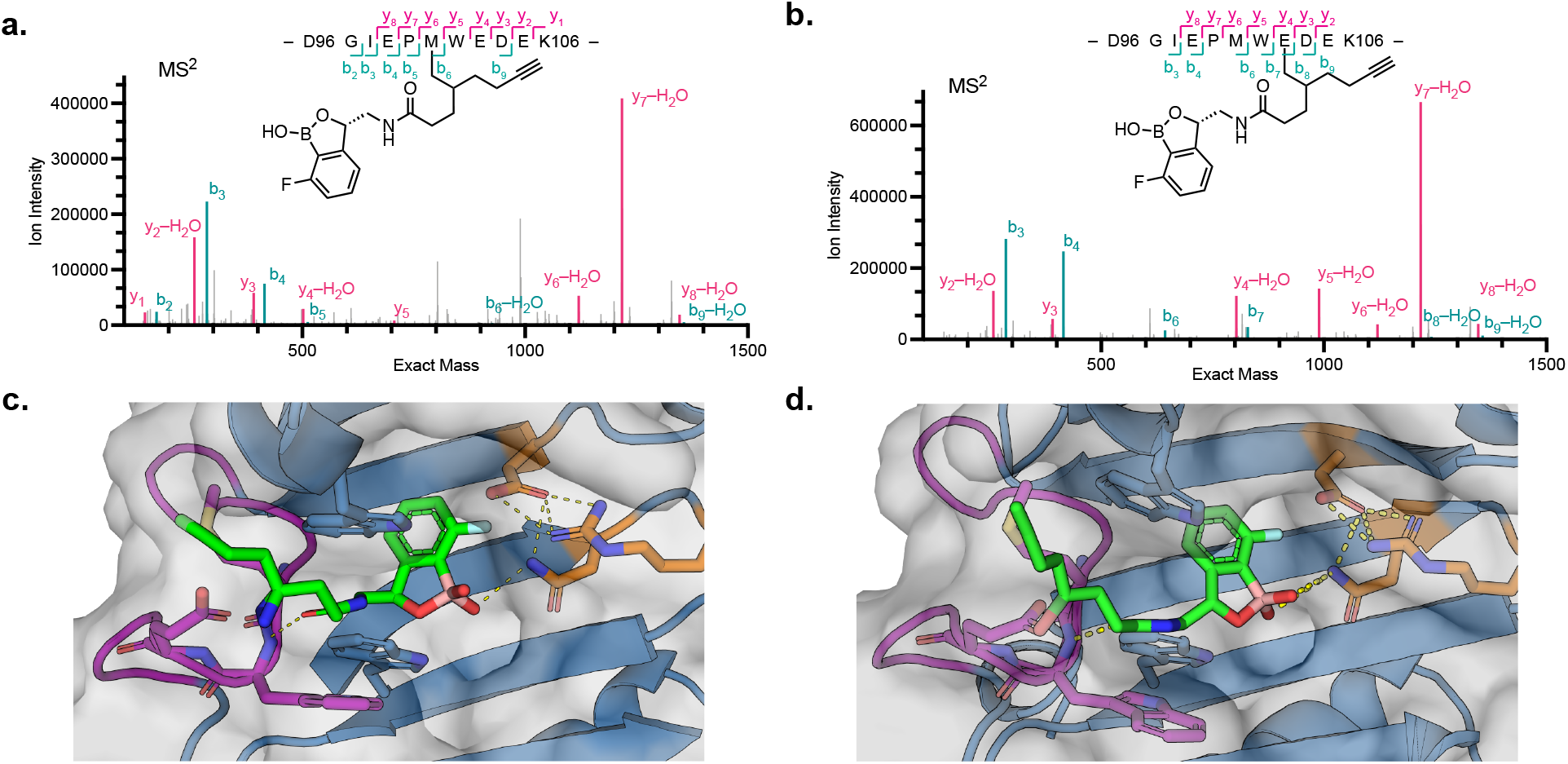
(a) MS^2^ of Met101 labeled peptide derived from *in vitro* labeling of eIF4E with (*S*)-DMP2. (b) MS^2^ of Glu103 labeled peptide from *in vitro* labeling of eIF4E with **(*S*)-DMP2**. (Duplicate) (c) Model of the anionic boronate form of **(*R*)-DMP2** reversibly bound to eIF4E. The covalently modified peptide is colored in purple and the individual residues are shown as sticks. Residues that play an important role in a hydrogen bonding network to the boronate are orange. Hydrogen bonds are indicated by dashed yellow lines. (AlphaFold 3) (d) Model of the anionic boronate form of **(*S*)-DMP2** covalently bound to eIF4E at Glu103. The scheme is consistent with panel C. (AlphaFold 3)

A model for the interactions that are driving the affinity of **(*S*)-DMP2** to eIF4E was developed using AlphaFold3 (AF3). AF3 successfully reproduced experimentally observed binding modes in four benchmark benzoxaborole bound crystal structures (Supplementary Figure 5).^39^ For modeling the eIF4E–**(*S*)-DMP2** complex we first reversibly docked the ligand into eIF4E but observed inversion of stereochemistry at the benzylic position yielding a model for eIF4E–**(*R*)-DMP2** (Figure 4c). This is a known problem for AF3, but the diazirine moiety was placed near both Met101 and Glu103.^39^ **(*R*)-DMP1** was also found to be enriched for labeling eIF4E in our chemoproteomics data (Figure 1c). By covalently ligating the diazirine bound carbon of **(*S*)-DMP2** to the carboxylic acid side chain of Glu103, AF3 produced models of the eIF4E–**(*S*)-DMP2** complex (Figure 4d). The model suggests that H-bonding occurs between the **(*S*)-DMP2** amide carbonyl and main chain amide N-H of Trp102 and the anionic boronate to the side chain of Asn155. These interactions were observed repeatedly across multiple models. The aromatic ring is also placed within a deep hydrophobic pocket that is not typically occupied when bound by m7GTP. Co-crystal structures of the acyclic Bn7GMP eIF4E ligands^26, 30^ also fill this pocket suggesting this mode of binding can help to drive eIF4E affinity.

In summary, a series of benzoxaboroles containing a diazirine/alkyne tag was synthesized to explore if this chemotype might reveal ligandability of challenging to drug targets. Benzoxaboroles substituted at the pro-chiral benzylic methylene had high enrichment for the proto-oncogene eIF4E while benzoxaboroles bearing substitutions at other positions displayed minimal to no enrichment. Additionally, enantiomeric pair probes revealed stereoselective protein binding preferences based on enrichment values and in-gel fluorescence bands for **(*S*)-DMP1** and **(*S*)-DMP2**. *In vitro* experiments with recombinant eIF4E revealed that the 5’ mRNA cap analogue **m7GTPG** competed with **(*S*)-DMP2** labeling of eIF4E. In cells, eIF4E labeled by **(*S*)-DMP1** or **(*S*)-DMP2** had diminished cap binding capabilities. Site of labeling experiments determined that **(*S*)-DMP2** modifies residues within eIF4E’s cap binding pocket. AF3 models generated from **(*S*)-DMP2** reversibly and covalently bound to eIF4E suggest that the amide bond connecting the PAL probe to the benzoxaborole scaffold makes important H-bonding contacts to the main chain of Trp102 while the anionic boronate H-bonds to the side chain of Asn155. The benzoxaborole scaffold of **DMP1** and **DMP2** is structurally unique compared to all other precedented cap competitive ligands for eIF4E while maintaining important contacts within the cap binding pocket.^24^

## Supporting information

Supplemental Proteomics Data

## ASSOCIATED CONTENT

### Supporting Information

The Supporting Information is available free of charge on the ACS Publications website.

Detailed experimental methods including synthesis and characterization of new compounds, protein expression and purification, chemoproteomics experimental set up and data analysis, and methods for all western blot and in gel fluorescence experiments. Raw chemoproteomics data can be accessed at https://massive.ucsd.edu/Prote-oSAFe/dataset.jsp?task=4f661123eaa14d4c932d296c74bbaf6e

## AUTHOR INFORMATION

### Funding Sources

J.B.C. was supported by a Ruth Kirschstein NRSA from the NCI training grant (T32CA108462).

D.M.P. was supported by a Ruth Kirschstein NRSA from the NCI of the NIH (F32CA253966).

G.B.C. was supported by the Tobacco-Related Disease Research Program Postdoctoral Fellowship Award (T32FT4880). National Institutes of Health grant U54CA243125.

### Notes

The authors declare the following competing financial interest(s): J.T. is a co-founder of Kezar Life Sciences and Terremoto Bio-sciences, and is a scientific advisor to Iambic Therapeutics.

K.M.S. has consulting agreements for the following companies, which involve monetary and/or stock compensation: BridGene Biosciences, Erasca, Exai, Genentech/Roche, Kumquat Biosciences, Kura Oncology, Lyterian, Merck, Montara, Nextech, Pfizer, Revolution Medicines, Rezo, Tahoe Therapeutics, Totus, Type6 Therapeutics, Wellspring Biosciences (Araxes Pharma).

D.M.P. is an employee of Novartis.

## ACKNOWLEDGMENT

We would like to thank the Sarpong Lab at UC Berkeley for generously allowing us to use their polarimeter to determine the optical rotation of our enantioenriched benzoxaboroles. We would also like to thank Hasan Celik at the UC Berkeley College of Chemistry NMR facility for his assistance in acquiring NMR spectra. We would also like to thank Lawrence Zhu for their assistance in expression and purification of truncated eIF4E, and Megan Moore for their early insights into immunoblotting for eIF4E.

---

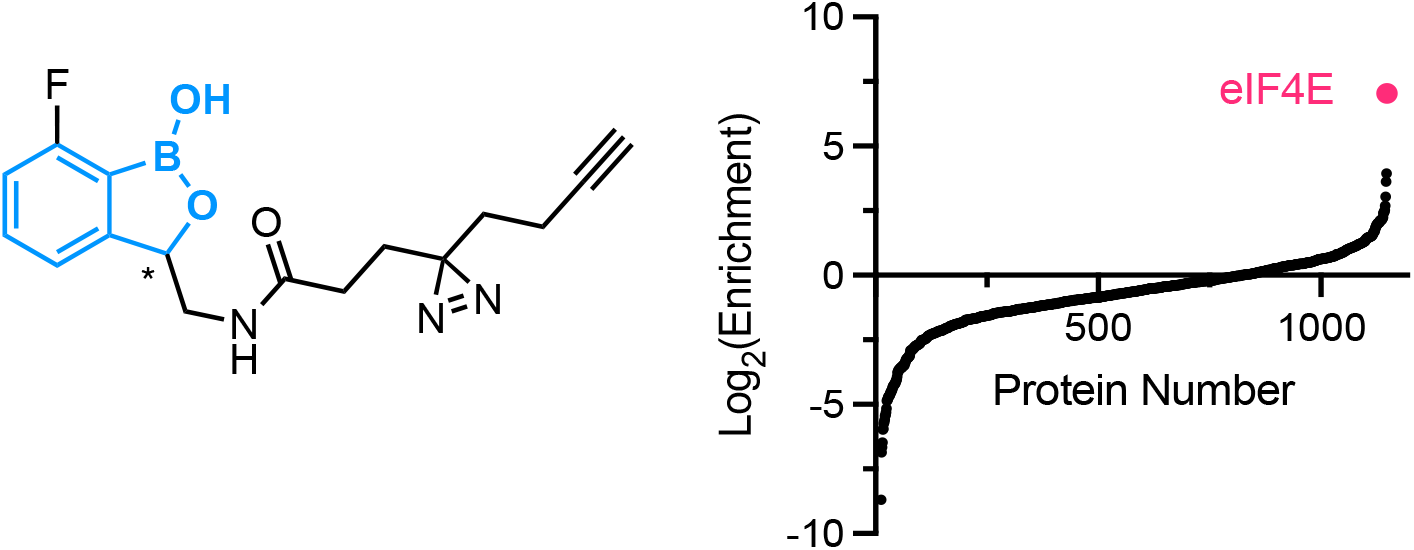

## Notes

https://doi.org/doi:10.25345/C5FX74B8H

### Summary of Updates

Additional Supplemental files figures were included to support the conclusions of the paper. The main body of the text was condensed, but the conclusions remain unchanged.

