## Supplementary Information for "Benzoxaboroles are structurally unique binders of eukaryotic translation initiation factor 4E"

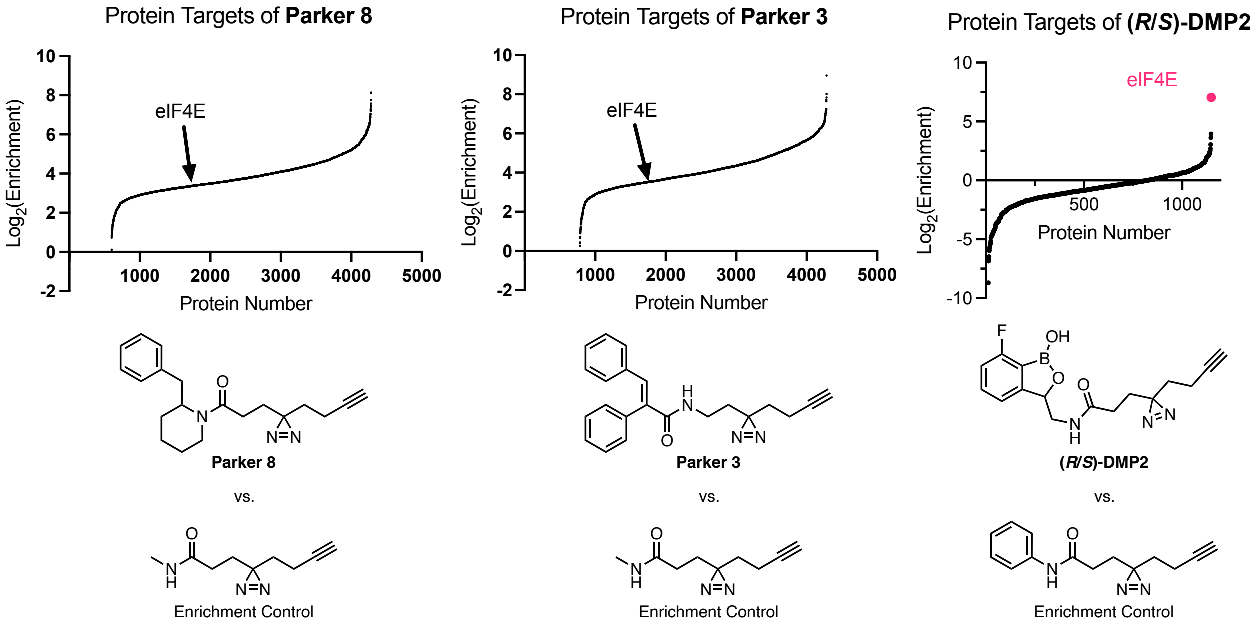

**Supplementary Figure 1.** Comparison of diazirine probes **Parker 8**, **Parker 3**, and **(*R*/*S*)-DMP2** that have been mapped to the cap binding pocket of eIF4E. The total enrichment values from Wozniak and co-workers were calculated from their reported enrichment values for each compound versus their methylamine derived enrichment control. All ligands were mapped to ligating the same eIF4E peptide (D96GIEPMWEDEK106).[1]

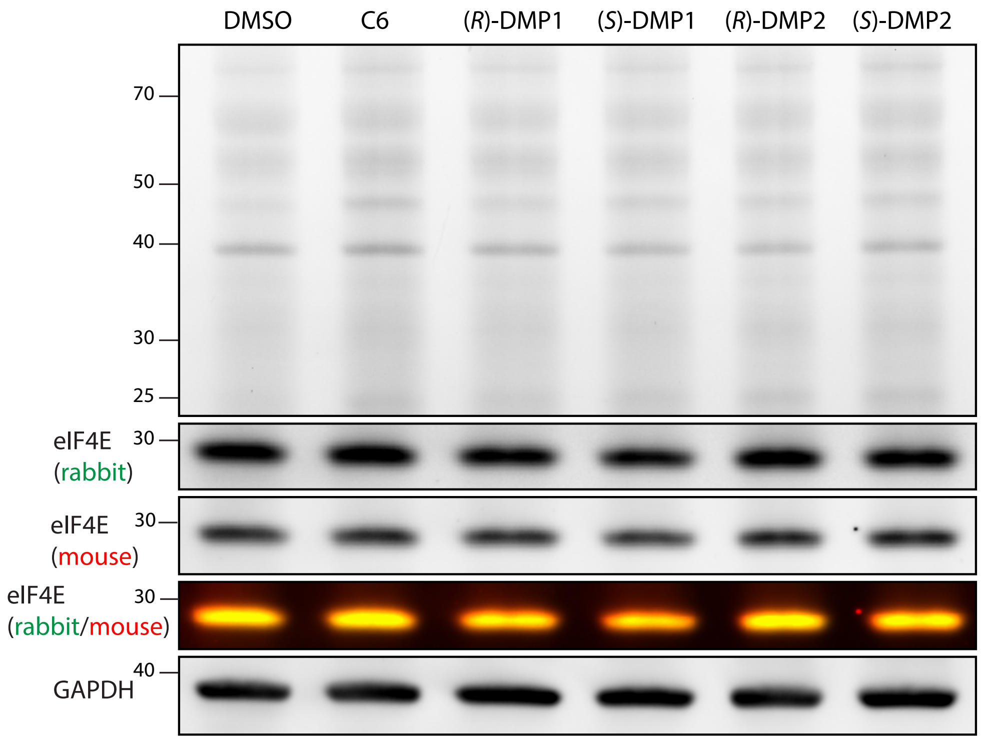

**Supplementary Figure 2.** HEK293T cells treated with 100 µM of the indicated compound or DMSO without UV exposure. The protocols for lysate generation and click reaction with TAMRA Plus Azide were identical to the procedures used in Figure 2a without the exposure to 365 nM light.

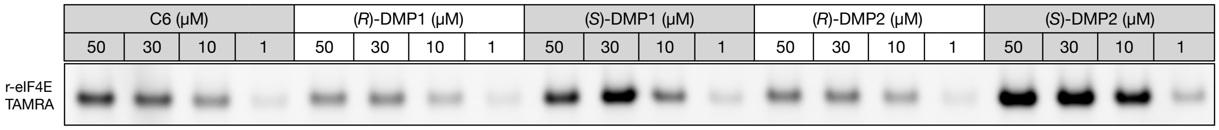

**Supplementary Figure 3.** Compound labeled recombinant eIF4E (28–217) visualized by in-gel fluorescence. (Duplicate)

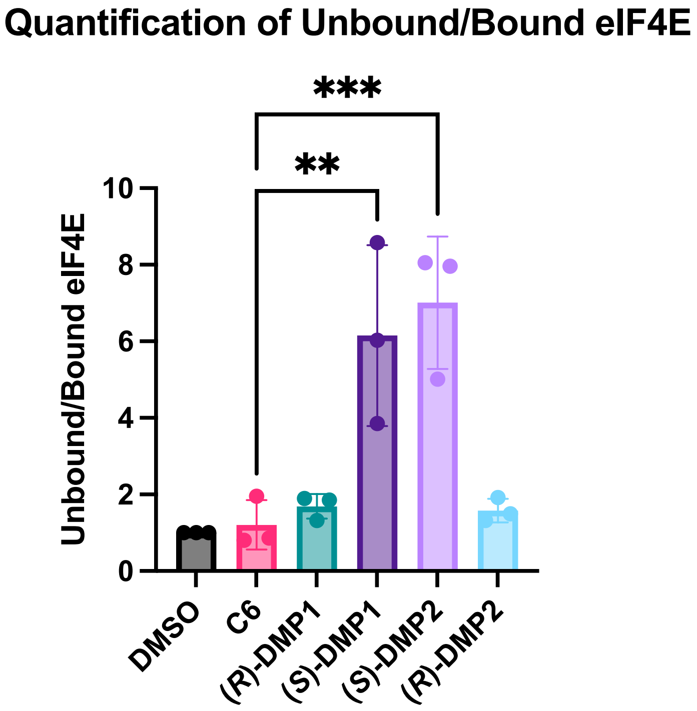

**Supplementary Figure 4.** Quantification of results in Figure 3b. Three independent chemoprecipitations of eIF4E from compound treated samples were quantified by western blot using the rabbit eIF4E primary antibody (Cell Signaling #2067). The adjusted signal (Image Lab) in the unbound fraction was DMSO corrected and divided by the adjusted signal in the DMSO corrected bound fraction. The adjusted P value for **(*S*)-DMP1** vs **C6** equals 0.0015 and **(*S*)-DMP2** vs **C6** equals 0.0004.

RMSD 2V0G

| Prediction Index | RMSD (A) |
|---|---|
| 0 | 0.2827359634004840 |
| 1 | 0.42865939275840100 |
| 2 | 0.3096634261581450 |
| 3 | 0.29160772023387900 |
| 4 | 0.3667620141181490 |
| Average | 0.3358857033338120 |

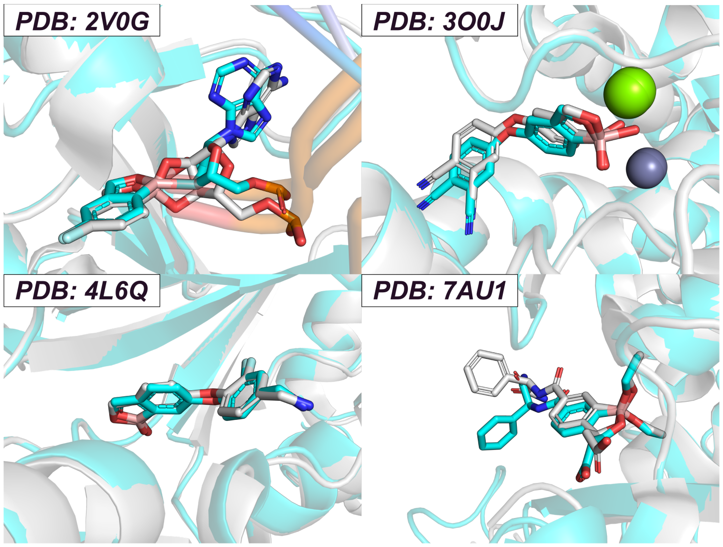

RMSD 3O0J

| Prediction Index | RMSD (A) |
|---|---|
| 0 | 0.6752678727734640 |
| 1 | 1.5767467245354700 |
| 2 | 0.7403121413746140 |
| 3 | 1.764478733224070 |
| 4 | 0.989347057407055 |
| Average | 1.1492305058629300 |

RMSD 4L6Q

| Prediction Index | RMSD (A) |
|---|---|
| 0 | 0.238793622 |
| 1 | 0.253389029 |
| 2 | 0.231707757 |
| 3 | 0.259053347 |
| 4 | 0.268532713 |
| Average | 0.250295294 |

RMSD 7AU1

| Prediction Index | RMSD (A) |
|---|---|
| 0 | 1.9541862161631000 |
| 1 | 1.7439230500945700 |
| 2 | 1.904357445195040 |
| 3 | 1.5468148368615100 |
| 4 | 1.5222350535116200 |
| Average | 1.7343033203651700 |

**Supplementary Figure 5.** Overlay of four published co-crystal structures containing benzoxaboroles and the AlphaFold 3 predicted structures.[2-5] The published crystal structures are colored in gray while the AlphaFold 3 structures are in blue. The calculated RMSD for all heavy atoms across the 5 different seeds and their average value are reported in the adjacent tables. Their values being below 2Å indicates good replication of the previously reported data.[6]

*Methods*

*Live-cell treatment and photo-affinity labeling*

**For TMT10plex chemoproteomic experiments:** HEK 293T cells (ATCC) were cultured in high glucose DMEM (Gibco 11995073) supplemented with 10% heat-inactivated FBS (Axenia Biologix). Cells were seeded into 10 cm dishes and grown to roughly 80% confluence. The media was aspirated, and the cells were gently washed with either DPBS (with Mg and Ca, Gibco 14040133) or supplement-free DMEM media. The media was replaced with media containing the indicated probe (100 µM) or DMSO (0.5% DMSO final, in all cases). The cells were incubated with the indicated probe for 30 min at 37 °C, then cooled on ice and irradiated with a 50 watt 365 nm lamp for 10 min. Photo-irradiated cells were harvested by scraping, centrifuging the suspension (500 g, 5 min, 4 °C), and aspirating the supernatant solution. Cell pellets were washed by resuspension in DPBS, centrifugation (500 g, 5 min, 4 °C), and removal of the supernatant twice more. The cell pellets were flash frozen with liquid $N_2$ and stored at –80 °C or thawed on ice for immediate use. The pellets were resuspended in 400 µL of pre-chilled lysis buffer, pipetted up and down 10 times, and incubated for 30 min on ice before clarification. The samples were centrifuged (21000 g, 20 min, 4 ºC) and the clear supernatant was transferred to fresh 1.5 mL microcentrifuge tubes. The total protein concentration of each sample was determined with a BCA assay versus BSA, and the final concentration of protein in each sample was adjusted to 1.5 mg/mL with the addition of more lysis buffer.

**For TAMRA in-gel fluorescence/western blot experiments**: HEK 293T cells (ATCC) were cultured in high glucose DMEM (Gibco 11995073) supplemented with 10% heat-inactivated FBS (Axenia Biologix). Cells were seeded into 6-well plates and grown to roughly 80% confluence. The media was aspirated, and the cells were gently washed with either DPBS or supplement-free DMEM media. For competition experiments with the covalent compound **Taunton 12**, the cells were pre-treated with 10 µM **Taunton 12** (0.1% DMSO final) or vehicle in DMEM + 10% FBS media and incubated at 37 ºC for 1 h. For non-competitive and competitive experiments, the media was replaced with DMEM media containing the indicated probe (5–100 µM) or DMSO (0.5% DMSO final, in all cases). The cells were incubated with the indicated probe for 30 min at 37 °C, then cooled on ice and irradiated with a 50 watt 365 nm lamp for 10 min. Media was aspirated from photo-irradiated cells, and the cells were gently washed with DPBS. The cells were then incubated with 150 µL of pre-chilled lysis buffer for 30 min, and the lysed samples were clarified by centrifugation (21000 g, 20 min, 4 ºC). The clear supernatant was transferred to fresh 1.5 mL microcentrifuge tubes. The total protein concentration of each sample was determined with a BCA assay versus BSA, and the final concentration of protein in each sample was adjusted to 1.0 mg/mL with the addition of more lysis buffer as needed.

*Lysis buffer*: 100 mM HEPES, 150 mM NaCl, pH 7.5 supplemented with NP-40 (0.1% v/v) and protease inhibitor (Roche cOmplete, EDTA-free, Sigma Aldrich 11836170001, 1 mini tablet per 10 ml)

*TMT-labeled cell sample preparation*

**Biotin "click"** A portion from each sample was diluted with buffer to obtain a 500 µl sample at 1.5 mg/ml total concentration. Freshly prepared Biotin "Click Mix" (55 µl) was added to each sample, and samples were mixed by rotator at ambient temperature for 1 h. Cold methanol (3 ml) was added to each sample and mixed, the proteins were pelleted by centrifugation (4700 g, 15 min, 4 °C), and the supernatant decanted. The pellets were washed by resuspending in 1:1 MeOH/$CHCl_3$ (1 ml) with bath sonication, centrifuging (4700 g, 15 min, 4 °C), and decanting the supernatant. The washing procedure was repeated again with 1:1 MeOH/$CHCl_3$ (1 ml) and finally with 3:1 MeOH/$CHCl_3$ (3 ml). The pellets were allowed to dry under air for approximately 3 min.

*Biotin "Click Mix:"* TBTA (92 µM in mix, 300 µl from 1.7 mM stock in 1:4 DMSO/*t*-BuOH), $CuSO_4$ (0.90 mM in mix, 100 µl from 50 mM stock in water), TCEP·HCl (0.90 mM in mix, 100 µl from 50 mM stock in water), and biotin PEG4 picolyl azide (90 µM in mix, 50 µl from 10 mM stock in DMSO, Sigma 900912)

**Streptavidin capture.** To each sample, urea (500 µl, 6 M in water) and SDS (10 µl, 10% w/v in DPBS) were added. The samples were homogenized with bath sonication, then a solution of TCEP and $K_2CO_3$ in DPBS (50 µl, 100 mM TCEP, 300 mM $K_2CO_3$, prepared immediately before adding) was added. The samples were incubated at 37 °C for 30 min. A solution of iodoacetamide in DPBS (70 µl, 400 mM) was added. The samples were incubated at room temperature in the dark for 30 min. A solution of SDS (130 µl, 10% w/v in DPBS) was added. The samples were diluted with DPBS (5.5 ml). To each sample, streptavidin-agarose beads (100 µl, 50% slurry, prewashed 3x with DPBS, Thermo 20353) were added. The samples were mixed by rotator overnight at 4 °C. An additional portion of streptavidin-agarose beads (50 µl) were added, and the samples were mixed by rotator at ambient temperature for an additional 90 min. The beads were collected by centrifugation (1400 g, 2 min, 18 °C) and decanting the supernatant. The beads were washed with SDS in DPBS (5 ml, 0.2% w/v, centrifuged 1400 g, 2 min, 18 °C), DPBS (twice, 5 ml, centrifuged 5000 g, 10 min, 4 °C), and EPPS (200 mM, pH 8, centrifuged 2000 g, 10 min, 4 °C).

**On-bead digestion.** Beads were resuspended in urea (2 M) in EPPS (200 µl, 200 mM, pH 8) and transferred to lo-bind tubes (1.5 ml, Eppendorf 022431021). $CaCl_2$ (1 µl, 0.25 M in water) and trypsin (2 µl, 1 mg/ml, Pierce 90057) were added. The samples were mixed on a shaker at 37 °C and 1000 rpm overnight (18 h). The samples were centrifuged (2000 g, 10 min, 4 °C), and the supernatant from each sample was carefully transferred to lo-bind tubes.

**TMT labelling.** To each sample, MeCN (94 µl, approximately 30% v/v final concentration) and the corresponding TMT reagent (6 µl of 20 µg $mL^{-1}$ stock, stocks reconstituted from kit with dry MeCN, TMT10plex kit, Thermo 90113) were added. The samples were mixed and incubated at ambient temperature for 1 h. Reactions were quenched by adding hydroxylamine (6 µl, 5% w/v), incubating for 5 min, and then adding formic acid (4 µl). Samples were frozen and concentrated on a centrifugal evaporator. To each sample, 92:5:3 $H_2O$/MeCN/FA was added, and samples were bath sonicated to obtain clear solutions. Samples were pooled in a clean lo-bind tube, rinsing the original tubes with additional portions of the same solvent mixture (final combined volume approximately 550 µl). The sample was mixed well then divided evenly into 5 lo-bind tubes. Peptides were captured with OMIX C18 tips (100 µl tip, Agilent A57003100, one tip per tube, preactivated with 3 × 100 µl MeCN + 0.1% formic acid followed by 3 × 100 µl $H_2O$ + 0.1% formic acid). Each tip was washed thrice with water (100 µl each), then peptides were eluted with 4:1 MeCN/$H_2O$ + 0.1% formic acid (3 × 100 µl). The resulting peptide solutions were frozen and lyophilized, then extracted with 95:5:0.1 $H_2O$/MeCN/formic acid (20 µl) and transferred to glass vials for LC-MS/MS analysis.

*LC-MS/MS injection and data analysis for whole cell lysate chemoproteomics*

Reconstituted TMT 10-plex samples were analyzed on an Orbitrap Eclipse Tribrid Mass Spectrometer (Thermo Fisher Scientific) connected to an Ultimate 3000 RSLCnano system with 0.1% formic acid in $H_2O$ as buffer A and 95% MeCN, 5% $H_2O$, 0.1% formic acid as buffer B. 5 µL of the samples were loaded for each injection (total of two injections per sample). Peptides were separated on an EASY-Spray 3 µm, 75 µm × 15 cm C18 column (Thermo Fisher Scientific, Cat #: ES800) with the following LC settings: flow rate at 0.3 µL $min^{-1}$, loading samples at 4% B for 20 min, then 4–7% B over 7 min, 7–35% B over 130 min, 35–95% B over 5 min, 95% B for 10 min, 95%–4% B over 0.5 min and finally 4% B for 9.5 min. MS2 and RTS-MS3 methods were utilized for TMT reporter quantification. In the MS2 method (one injection), MS1 spectra were acquired at a resolution of 120 K with m/z scan range of 400–1600, RF lens of 30%, a maximum ion injection time of 50 ms, charge states of 2–6, and a 1 scan 60s dynamic exclusion time. MS2 spectra were acquired via HCD at a collision energy of 40%, in the orbitrap with an isolation width of 0.5 m/z, a resolution of 50 K, m/z scan range of 100–3000 and a maximum ion injection time of 100 ms. In the RTS-MS3 method (one injection), the MS1 settings were the same as in the MS2 based method. For the RTS-MS3 method, MS2 spectra were acquired via CID at a collision energy of 35%, in the ion trap with an isolation width of 0.5 m/z, and m/z scan range of 400–1600. For the real time search, which is based on the Comet search engine, MS2 spectra were searched against the HUMAN reviewed Swiss-Prot FASTA database with the digestion enzyme set to trypsin. Methionine oxidation was set as a variable modification, while carbamidomethylation of cysteine and TMT modification of the N-terminal amine and Lys were set as static modifications. For MS3 acquisition, a synchronous precursor selection (SPS) of 10 fragments was acquired in the orbitrap for a maximum injection time set to Auto. MS3 spectra were collected at a resolution of 60 K using the orbitrap with the HCD collision energy set at 55%.

The LC-MS/MS data were searched using MaxQuant (v.1.6.7.0) against the HUMAN reviewed Swiss-Prot FASTA database. Under "Group-specific parameters", for MS2 data, the "type" was set as "Reporter ion MS2". For RTS data, the "type" was set as "Reporter ion MS3". The "Isobaric labels" was set as "10plex TMT" and the reporter ion isotopic distributions were incorporated to correct for impurities during synthesis of the TMT reagents according to the manufacturer's specifications. Methionine oxidation and protein N-terminal acetylation were set as variable modifications, while carbamidomethylation of cysteine was set as a static modification. "Trypsin/P" was selected as the digestion enzyme with a maximum of 2 missed cleavages. All other parameters were set as default. TMT intensities in each channel were normalized such that the median TMT intensity values (based on all modified sites) were equivalent across all channels. Median TMT intensity values for each benzoxaborole were then divided by the respective TMT intensity value from control **C6** to calculate the reported enrichment values.

*In-gel fluorescence and western blot experiments from cell lysate*

3 µL of TAMRA "Click Mix" was added to 22 µL of each treated cell lysate with a corrected protein concentration of 1 mg $mL^{-1}$. The click reaction was incubated in the dark at 23 ºC for 90 min before quenching the reaction with 7 µL of 4× SDS loading buffer (BioRad 1610747). The quenched mixture was vortexed and left to denature at 23 ºC for 15 min. Each sample was then loaded (10–15 µL) into a 4–12% bis-tris protein gel (ThermoFisher NP0321BOX). The gel was run in MES running buffer (Fisher Scientific NP000202) at 200 V for 42 min, or until the loading/unreacted TAMRA dye reached the bottom of the gel. The gel was briefly rinsed with water, and then imaged on BioRad's ChemiDoc instrument at 550 nm. The gel was then transferred onto a nitrocellulose membrane (ThermoFisher IB33001X3) using an iBlot 3 Western Blot dry transfer system on the medium sized protein transfer setting. The membranes were then cut into the desired strips and blocked with 5% w/v BSA/TBST (BSA: Sigma-Aldrich 12659; TBST: 20 mM Tris, 150 mM NaCl, 0.1% Tween-20, plus 0.02% sodium azide) blocking buffer for 1 h at 23 ºC. The membrane was then incubated with blocking buffer containing a 1:1000 dilution of rabbit eIF4E (Cell Signaling #2067) primary antibody and GAPDH (Proteintech 60004-1-Ig) primary antibody and 1:500 dilution of the mouse eIF4E (BD Biosciences Cat #610269) primary antibody overnight (16 h) at 4 ºC. The same results were observed if all the antibodies were multiplexed or run individually. The primary bound membrane was washed with 1× TBST (20 mM Tris, 150 mM NaCl, 0.1% Tween-20, plus 0.02% sodium azide) for 3 × 5 min. The rinsed membrane was then incubated with a 1:5000 dilution of LI-COR IRDye 800CW goat anti-rabbit (926-32211) and IRDye 680RD goat anti-mouse (926-68070) antibodies in 5% w/v BSA/TBST for 1 h at 23 ºC. The secondary bound membrane was again washed with 1× TBST for 3 × 5 minutes. The prepared membrane was then imaged on BioRad's ChemiDoc instrument on the 800 and 680 channels with the automatic sensitivity setting. The scans were processed in ImageJ or Image Lab software. Each raw image was cropped into individual strips, and brightness of each strip was adjusted independently.

*TAMRA "Click Mix"*: 33 µL of 1.7 mM TBTA in 1:4 DMSO/*t*-BuOH, 11 µL 50 mM $CuSO_4$ in $H_2O$, 11 µL 50 mM TCEP in $H_2O$, and 11 uL of 1.25 mM TAMRA Azide Plus (BroadPharm BP-40173)

The rabbit eIF4E (Cell Signaling #2067) primary antibody is monoclonal with no reported epitope. The mouse eIF4E (BD Biosciences Cat #610269) primary antibody is monoclonal with no reported epitope.

*Recombinant eIF4E (G–28–217) expression and purification*

A plasmid encoding for full length eIF4E with an N-terminal 6×His tag followed by a TEV cleavage site under the control of a Lac promoter and an ampicillin resistance gene was purchased from Genscript. The base pairs encoding for the first 27 amino acids were truncated from the plasmid using PCR amplification (NEB E0554) and blunt end ligation (NEB M0554S). BL21(DE3) *E. coli* were transformed with the truncated plasmid and plated directly onto LB agar plates containing 50 µg/mL carbenicillin. The plate was incubated at 37 ºC for 16 h, and then stored at 4 ºC until ready to inoculate the starter culture.

10–20 colonies were added to 200 mL of LB supplemented with carbenicillin and incubated in a 37 ºC shaker overnight (16 h). 25 mL of the starter culture was used to inoculate 1 L of TB supplemented with carbenicillin, which was then placed in a 37 ºC shaker until the optical density ($OD_{600}$) reached 0.6. The culture was then induced with IPTG (0.2 mM final) and placed in an 18 ºC shaker for 18 h. The culture was then transferred to a 1 L centrifuge tube and pelleted (4000 RPM, 15 min). The supernatant was decanted, and the cell pellet was transferred to a 50 mL falcon tube. The tube was flash frozen in liquid $N_2$ and stored at –80 ºC until ready for lysis and purification.

The cell pellet was thawed on ice and placed in a pre-cooled metal cup. The cells were suspended in 30 mL lysis buffer (50 mM HEPES pH 7.5, 150 mM NaCl) supplemented with EDTA-free protease inhibitor and lysed by ultrasonication. The lysate was then clarified by centrifugation for 1 h at 16000 g and 4 ºC. The lysate was collected and placed into a 50 mL falcon tube on ice.

The clarified lysate was purified by immobilized metal affinity chromatography (IMAC) at 4 ºC. The His-TEV tagged protein was captured on Ni resin (HisPur™ Ni-NTA Resin from Thermo Scientific 88221) for 2 h with agitation of the supernatant and beads every 15 minutes followed by elution of the flow through. The beads were then washed with 2 × 8 mL of wash buffer (50 mM HEPES pH 7.5, 150 mM NaCl, 5 mM imidazole), 2 × 8 mL of high salt buffer (50 mM HEPES pH 7.5, 1 M NaCl, 5 mM imidazole), and 8 mL of wash buffer. The non-specific bound proteins were eluted with 8 mL of high imidazole wash buffer (50 mM HEPES pH 7.5, 150 mM NaCl, 50 mM imidazole) and 8 mL of a 1:1 mixture of high imidazole wash buffer and elution buffer (50 mM HEPES pH 7.5, 150 mM NaCl, 200 mM imidazole). The remaining bound proteins were eluted with 4 mL aliquots of elution buffer. A colormetric protein concentration assay (Bradford) was used to determine the end of the protein elution.

His-tagged TEV protease was added to the pooled protein fractions, which was then placed into a 70 mL <10K MWCO dialysis cassette. The mixture was dialyzed against TEV cleavage buffer (30 mM HEPES pH 7.4, 150 mM NaCl, 10 mM glycerol, 5 mM BME, 0.2 mM GDP) at 4 ºC until complete cleavage of the TEV site was observed by LC-MS (16 h).

The cleaved protein was then captured on Ni resin at 4 ºC for 2 h. The column was then washed with 8 mL of wash buffer and 3 × 8 mL of high imidazole wash buffer. The remaining eIF4E on the column was then eluted with 8 mL fractions of 2:1 high imidazole wash buffer to elution buffer. Collected fractions were run on a 4–12% bis-tris gel in MES buffer to determine which fractions would be carried forward for size exclusion chromatography (SEC).

The pooled fractions were purified by SEC using a Superdex 75 10/300 GL column (GE Healthcare Life Sciences) with SEC buffer (30 mM HEPES pH 7.5, 100 mM NaCl, 10 mM $MgCl_2$), and the pure fractions were again pooled, concentrated, flash frozen in liquid $N_2$, and stored at –80 ºC.

*In vitro recombinant eIF4E (G–28–217) labeling experiments*

**Recombinant protein labeling.** All labeling experiments were run in a final volume of 40 µL(<0.5% DMSO) in clear bottomed 96 well plates, and all reaction plates were prepared on ice. The requisite amount of SEC buffer (30 mM HEPES pH 7.5, 100 mM NaCl, 10 mM $MgCl_2$) to bring each reaction to a final volume of 40 µL was added to each well. Recombinant eIF4E (G–28–217) was then added to each well to reach a final concentration of 10 µM. **Taunton 12**, m7GTPG, and GTPG were added to their corresponding wells to reach final concentrations of 100 µM, 10 µM, and 10 µM respectively. A 20 mM stock solution of the indicated compound in DMSO was diluted 1:100 in SEC buffer and was then added to each well to reach final concentrations of 50, 30, 10, or 1 µM and incubated for 1 h in the dark on ice. The plates were then photoirradiated with 365 nm light for 10 min on ice.

**In-gel fluorescence.** 22 µL of each reaction on recombinant eIF4E was placed into a PCR tube and mixed with 3 µL of TAMRA "Click Mix" as described in *In-gel fluorescence and western blot experiments from cell* lysate. The click reaction was placed in a dark drawer at 23 ºC for 90 min before quenching the reaction with 8 µL of 4× SDS loading buffer (BioRad 1610747). The quenched mixture was vortexed and left to denature at 23 ºC for 15 min. Each sample was then loaded into a 4–12% bis-tris protein gel from Invitrogen. The gel was run in MES running buffer at 200V for 42 min or until the loading/unreacted TAMRA dye reached the bottom of the gel. The gel was then briefly rinsed with DI water and imaged on BioRad's ChemiDoc instrument at 550 nm to visualize the TAMRA labeled protein.

*m7GTP agarose bead cell lysate pull down and quantification*

For each compound treatment tested, 200 µL of γ-aminophenyl-$m^7$GTP ($C_{10}$-spacer)-agarose beads (Jena Bioscience AC-155L) were pipetted into a spin column (Thermo Scientific PI69725). The supernatant was removed with a benchtop centrifuge and the beads were washed with 2 × 500 µL equilibration buffer (100 mM HEPES pH 7.5, 150 mM NaCl, 0.1% NP-40 v/v). 1 mg/mL compound treated cell lysate was further diluted with lysis buffer to 0.5 mg/mL and 80 µL of the diluted compound treated lysate was added onto the beads and incubated for 1.5 h at 4 ºC on a shaker. The incubated lysate was then collected as the unbound fraction by centrifugation into a 1.5 mL microcentrifuge tube. The protein bound beads were then washed with 5 × 500 µL equilibration buffer and then resuspended in 80 µL equilibration buffer. The resuspended protein bound beads were then diluted with 4× SDS loading buffer and denatured at 95 ºC for 5 min. The supernatant on the beads was immediately collected in clean 1.5 mL microcentrifuge tubes. The input cell lysates and the unbound protein fractions were diluted with 4× SDS loading buffer and denatured at 95 ºC for 5 min. Each sample was loaded into a 4–12% bis-tris gel, the gel was ran in MES, and then transferred and analyzed by western blot. The protocols for running the gel, transfer, and western are identical to those found in *In-gel fluorescence and western blot experiments from cell lysate*.

The western bands corresponding to the eIF4E rabbit antibody signals were quantified from 3 unique replicates using Image Lab. The adjusted volume value, as reported by Image Lab, was normalized to DMSO within a given replicate for each sample (Ex. DMSO normalized signal for the unbound fraction of a sample treated with (*S*)-DMP2 = Unbound rabbit signal for (*S*)-DMP2/unbound rabbit signal for DMSO). Within a given western experiment, the DMSO normalized eIF4E signal values for the unbound fraction were divided by the normalized bound value to give the ratio of Unbound/Bound eIF4E signal for each compound:

$$\frac{Unbound}{Bound}\ ratio = \left[\frac{eIF4E\ signal\ in\ unbound\ treated\ lane}{eIF4E\ signal\ in\ unbound\ DMSO\ lane}\right] / \left[\frac{eIF4E\ signal\ in\ bound\ treated\ lane}{eIF4E\ signal\ in\ bound\ DMSO\ lane}\right]$$

The DMSO normalized Unbound/Bound eIF4E values for each compound were plotted into a bar graph using PRISM, and statistical significance was measured using an ordinary one-way ANOVA.

*Site of labeling on recombinant protein sample preparation and LC-MS/MS method*

Recombinant eIF4E (G–28–217) was labeled with 50 mM **(*S*)-DMP2** following the protocol from the *In vitro recombinant eIF4E (G–28–217) labeling experiments*: **Recombinant protein labeling** section. The 40 µL mixture of labeled protein was diluted with 40 µL of 20 mM Tris (pH 8.0), 2 mM $CaCl_2$ buffer. The solution was sequentially incubated with DTT (1 µL, 400 mM, 50 ˚C, 30 min), iodoacetamide (4 µL, 200 mM, room temperature, 15 min), DTT (2.1 µL, 400 mM. room temperature, 15 min), and then mass spectrometry grade trypsin (Promega, Cat #: VA9000) (5 µL, 0.5 mg $mL^{-1}$, 37 ˚C, 18 h). The resulting peptides were acidified with 2 µL formic acid, desalted with C18 Omix Tips (Agilent, Cat#: A57003100) and eluted with 50% MeCN, 0.1% formic acid. The samples were dried down by SpeedVac and then analyzed by LC-MS/MS.

Tryptic peptides were reconstituted in 100 µL of 0.1% trifluoroacetic acid in water and were analyzed on an Orbitrap Eclipse Tribrid Mass Spectrometer (Thermo Fisher Scientific) connected to an Ultimate 3000 RSLCnano system with 0.1% formic acid in $H_2O$ as buffer A and 95% MeCN, 5% $H_2O$, 0.1% formic acid as buffer B. 5 µL of the samples were loaded for each injection (total of two injections per sample). Peptides were separated on an EASY-Spray 3 µm, 75 µm x 15 cm C18 column (Thermo Fisher Scientific, Cat #: ES800) with the following LC settings: variable flow rate at 0.3 µL $min^{-1}$ or 0.6 µL $min^{-1}$, loading samples at 2% B (0.6 µL $min^{-1}$) for 12 min, then 2% B (0.3 µL $min^{-1}$) over 0.1 min, then 2–31.6% B (0.3 µL $min^{-1}$) over 71.9 min, 31.6–52.6% B (0.3 µL $min^{-1}$) over 2 min, 52.6–2% B (0.3 µL $min^{-1}$) over 2 min, and finally 2% B (0.6 µL $min^{-1}$) for 7 min. In the static MS2 method (one injection), MS1 spectra were acquired at a resolution of 120 K with m/z scan range of 375–1500, RF lens of 30%, a maximum ion injection time of 50 ms, charge states of 2–7, and a 1 scan 30s dynamic exclusion time. MS2 spectra were acquired via HCD at a collision energy of 30%, in the orbitrap with an isolation width of 1.6 m/z, a resolution of 30 K, m/z scan range set to Auto and a maximum ion injection time of 100 ms. In the stepwise MS2 method (one injection), MS1 spectra were acquired at a resolution of 120 K with m/z scan range of 200–1500, RF lens of 30%, a maximum ion injection time of 50 ms, charge states of 2–7, and a 1 scan 30s dynamic exclusion time. MS2 spectra were acquired via HCD at a variable collision energy of 20%, 25%, and 30% in the orbitrap with an isolation width of 1.6 m/z, a resolution of 30 K, m/z scan range set to Auto and a maximum ion injection time of 100 ms.

The LC-MS/MS data were searched using MaxQuant (v.2.6.5.0) against an eIF4E FASTA. Under "Group-specific parameters", the "type" was set as "Standard". Methionine oxidation and **DMP2** modification on E, M, P, W, D, G, I, or K were set as variable modifications, while carbamidomethylation of cysteine was set as a static modification with the max number of modifications per peptide set to 5. "Trypsin/P" was selected as the digestion enzyme with a maximum of 2 missed cleavages. Under the "Protein quantification" section of "Global parameters", methionine oxidation and **DMP2** modification on E, M, P, W, D, G, I, or K were turned on for protein quantification. All other parameters were set as default.

*RMSD calculation*

Pocket-aligned RMSDs were computed following the procedure described in the AlphaFold3 Supporting Information.[6] Briefly, the binding pocket is defined as all heavy-atom coordinates located within 10 Å of any heavy atom of the ligand in the reference crystal structure. The predicted model is then superposed onto the ground-truth structure by least-squares rigid-body alignment using only the Cα atoms of residues within this pocket. Finally, the RMSD is calculated over all heavy atoms of the ligand.

*Chemical synthesis*

*General experimental details.* Solvents referred to as "dry" were either purchased (Acros extra dry) or dried with a solvent system containing activated molecular sieves. Other solvents were purchased from Fisher Scientific or Sigma Aldrich and used as received (ACS reagent or HPLC grade). Thin layer chromatography (TLC) was performed with indicator-containing glass-backed silica plates (F254, EMD) and visualized under a 254 nm UV lamp. Liquid chromatography mass spectroscopy (LC/MS) was performed with a Waters Acquity UPLC and Xevo QTOF. Normal-phase flash column chromatography was performed on a Teledyne Isco Combiflash Rf+ system. Reverse-phase (C18) flash column chromatography was performed on a Teledyne Isco EZPREP system. Preparative reverse-phase high pressure liquid chromatography (HPLC) was performed on either a Waters Autopurification or Teledyne Isco EZPREP system equipped with a 20x300 C18 column and UV/vis detector. NMR spectra were acquired on a 400 MHz Bruker Avance series instrument at the UCSF Pharm NMR lab or on a 600 MHz Bruker Avance series instrument at the UC Berkeley College of Chemistry NMR facility. $^{1}$H and $^{13}$C chemical shifts were referenced to the solvent peak ($CHCl_3$ $^{1}$H δ 7.26 ppm, $CDCl_3$ $^{13}$C δ 77.16, DMSO-*d*$_5$ $^{1}$H δ 2.50 ppm, DMSO-*d*$_6$ $^{13}$C δ 39.52). $^{19}$F NMR chemical shifts were referenced indirectly with Ξ = 94.094011.

*Precautions taken to avoid inadvertent photolysis of diazirine-containing compounds.* Reactions were either performed in amber vials or in clear vials wrapped in aluminum foil. Workups and other manipulations were performed with the hood lights off and away from light sources. The UV/vis detectors of Teledyne Isco systems were disabled during purification. Samples were stored in amber vials at –20 °C.

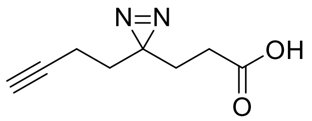

**3-(3-(But-3-yn-1-yl)-3*H*-diazirin-3-yl)propanoic acid.**

Prepared according to the procedures published by Parker et al.[7] $^{1}$H NMR (400 MHz, $CDCl_3$) δ 2.18 (t, *J* = 7.7 Hz, 2H), 2.06 – 1.98 (m, 3H), 1.82 (t, *J* = 7.7 Hz, 2H), 1.66 (t, *J* = 7.3 Hz, 2H). *Note: the acidic -$CO_2H$ peak was not observed in the $^{1}$H NMR spectrum.*

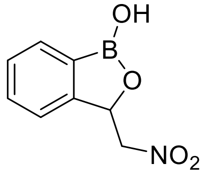

**(*rac*)-3-(Nitromethyl)benzo[*c*][1,2]oxaborol-1(3*H*)-ol** was prepared as previously reported.[8]

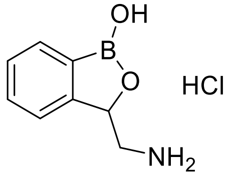

**(*rac*)-3-(Aminomethyl)benzo[*c*][1,2]oxaborol-1(3*H*)-ol hydrochloride.**

In a 100 ml round-bottom flask under Ar, Raney Nickel (4 ml, 50% aqueous slurry, approximately 34 mmol, 3.4 equiv) was suspended in MeOH (24 ml) and cooled to 0 °C in an ice bath. Methanolic ammonia (7 M, 12 ml, 84 mmol, 8.4 equiv) was added with stirring (total volume approximately 40 ml, final [$NH_3$] = 2 M). 3-

(Nitromethyl)benzo[*c*][1,2]oxaborol-1(3*H*)-ol (1.93 g, 10.0 mmol) was added to the reaction flask with stirring. The atmosphere was purged with hydrogen, and the mixture was stirred at 0 °C under an atmosphere of hydrogen for 8 h. The atmosphere was purged with Ar. The solution was filtered through a pad of Celite, which was rinsed thrice with MeOH (10 ml each). Volatiles were removed at reduced pressure. The residue was suspended in dry dioxane (30 ml). HCl (4 M in dioxane, 10 ml, 40 mmol, 4 equiv) was added with stirring. The solid precipitate was collected by filtration and dried under vacuum to give the crude racemate of the title compound. This sample was used in the next step without further purification.

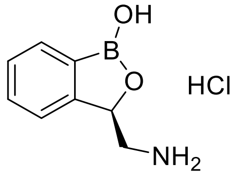

**(*R*)-3-(Aminomethyl)benzo[*c*][1,2]oxaborol-1(3*H*)-ol hydrochloride**.

The crude racemic 3-(aminomethyl)-benzo[*c*][1,2]oxaborol-1(3*H*)-ol hydrochloride obtained as described above was suspended in MeOH (10 ml). Sodium hydroxide (0.40 g, 10 mmol, 1.0 equiv) in MeOH (10 ml) was added with stirring, and the mixture was stirred at ambient temperature for 20 min. The mixture was filtered through a pad of Celite, which was rinsed thrice with MeOH (10 ml each), to give a clear blue solution. (*R*)-Mandelic acid ((*R*)-MA) (1.52 g, 10.0 mmol, 1.0 equiv) was added with stirring, and the mixture was stirred at ambient temperature for 20 min. This precipitate was collected by filtration, rinsed thrice with EtOH (5 ml each), and dried under vacuum to give the (*R*)-MA salt of title compound as a white powder (1.2 g, 2.8 mmol). Marfey analysis determined that this sample contained the (*R*) enantiomer in 73% ee. The filtrate was saved (see below).

This sample was recrystallized by dissolving in boiling EtOH (300 ml), filtering through a pad of Celite, cooling to 4 °C, and letting sit for 2 d. The crystals were collected by filtration and dried under vacuum to give the (*R*)-MA salt of title compound. This sample was determined to be >98% ee (*R*) by Marfey analysis. This sample was stirred in a solution of HCl (4 M in dioxane, 10 ml) at ambient temperature for 30 min, then diluted with diethyl ether (50 ml) and stirred for an additional 10 min. The solid was collected by filtration to give the title compound as a white powder (398 mg, 1.99 mmol, 19% yield).

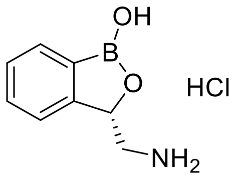

**(*S*)-3-(Aminomethyl)benzo[*c*][1,2]oxaborol-1(3*H*)-ol hydrochloride**.

The saved filtrate from the first step of the previous compound was concentrated then suspended in a solution of HCl (4 M in dioxane, 10 ml). The mixture was stirred at ambient temperature for 30 min then diluted with diethyl ether (50 ml). The solid was collected by filtration and rinsed thrice with ether (15 ml each) to obtain an off-white powder (1.17 g). This sample was dissolved in MeOH (5 ml). Sodium hydroxide (0.20 g, 5.0 mmol, 0.50 equiv) in MeOH (5 ml) was added with stirring, and the mixture was stirred at ambient temperature for 20 min. The mixture was concentrated then redissolved in absolute ethanol (30 ml). The mixture was filtered through a pad of Celite, which was rinsed thrice with EtOH (10 ml each). (*S*)-Mandelic acid ((*S*)-MA) (761 mg, 5.01 mmol, 0.50 equiv) was added with stirring, and the mixture was stirred at ambient temperature for 15 min. This precipitate was collected by filtration and rinsed twice with EtOH (10 ml each).

This sample was recrystallized by dissolving in boiling EtOH (340 ml), filtering through a pad of Celite, cooling to 4 °C, and letting sit for 2 d. The crystals were collected by filtration and dried under vacuum to give the (*S*)-MA salt of title compound. This sample was determined to be >98% ee (*S*) by Marfey analysis. This sample was stirred in a solution of HCl (4 M in dioxane, 10 ml) at ambient temperature for 30 min, then diluted with diethyl ether (50 ml) and stirred for an additional 10 min. The solid was collected by filtration to give the title compound as a white powder (316 mg, 1.58 mmol, 16% yield).

**Procedure for Marfey analysis.** In a 1-dram vial, the sample (either HCl or Mandelic acid salt form, approximately 1 mg, 4 umol) was dissolved in a solution of 1-fluoro-2-4-dinitrophenyl-5-L-alanine amide (FDAA) (0.35 ml, 10 mg/ml, 13 µmol, approximately 3 equiv) in acetone. Aqueous $NaHCO_3$ (50 µl, 1.0 M, 50 µmol, approximately 10 equiv) was added with stirring. The mixture was allowed to react at ambient temperature for 90 min. An aliquot of the suspension (20 µl) was removed and diluted with 1:1 MeCN/$H_2O$ with 0.1% FA (80 µl) to obtain a clear yellow solution, which was analyzed by analytical LC/MS (Waters UPLC, C18, 20-30% MeCN gradient in water with 0.1% formic acid). Example chromatograms are shown below.

Absolute configuration was assigned based on previous reports.[8, 9]

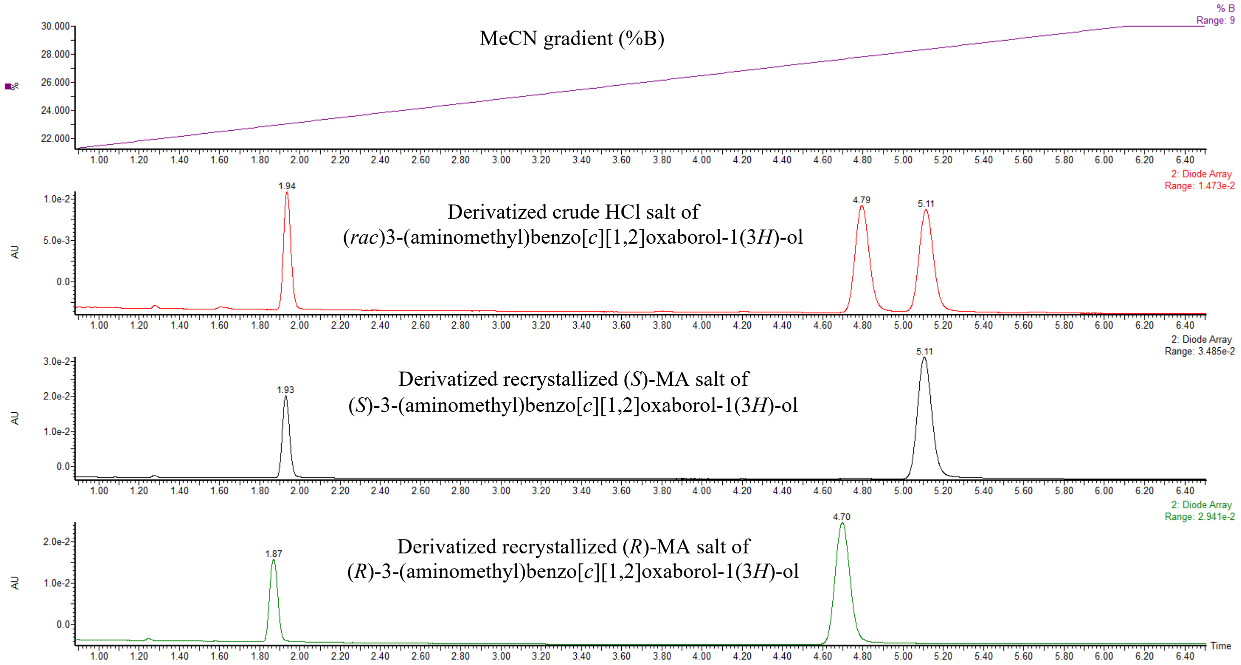

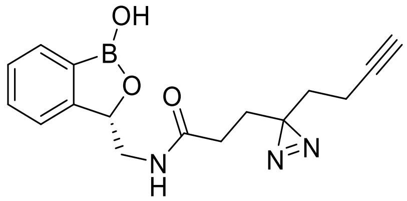

**(*S*)-3-(3-(But-3-yn-1-yl)-3*H*-diazirin-3-yl)-*N*-((1-hydroxy-1,3-dihydrobenzo[*c*][1,2]oxaborol-3-yl)methyl)propanamide ((*S*)-DMP1).**

An amber 1-dram vial under Ar was charged with 3-(3-(but-3-yn-1-yl)-3*H*-diazirin-3-yl)propanoic acid (83.2 mg, 0.500 mmol), (*S*)-3-(aminomethyl)-benzo[*c*][1,2]oxaborol-1(3*H*)-ol hydrochloride (110 mg, 0.552 mmol, 1.10 equiv), and HATU (285 mg, 0.750 mmol, 1.5 equiv). The mixture was cooled to 0 °C in an ice-water bath. Dry DMF (2.0 ml) and DIPEA (0.26 ml, 0.19 g, 1.5 mmol, 3.0 equiv) were added in that order with stirring. After 30 min, the cooling bath was removed, and the mixture was stirred at ambient temperature overnight. The mixture was diluted with EA (15 ml) and washed twice with $HCl_{(aq)}$ (1.0 M) and once with brine (15 ml each). The aqueous layers were extracted twice further with EA (20 ml each) in the same sequence. The organic layers were dried over $Na_2SO_4$ and filtered, and volatiles were removed at reduced pressure. The crude product was purified by flash column chromatography (MeOH gradient in DCM, then repurified with EA gradient in Hex, $R_f$ = 0.21 in 50% EA/Hex) followed by trituration with 1:1 ether/Hex (3 ml) at 4 °C. The solid was collected by vacuum filtration and washed twice with cold hexanes (2 ml each) to give the title compound as a white powder (79.2 mg, 0.255 mmol, 51% yield). $^1$H NMR (600 MHz, DMSO-*d*$_6$) δ 9.21 (s, 1H), 8.09 (t, *J* = 5.5, 4.8 Hz, 1H), 7.72 (d, *J* = 7.2 Hz, 1H), 7.46 (t, *J* = 7.2 Hz, 1H), 7.40 (d, *J* = 7.6 Hz, 1H), 7.35 (t, *J* = 7.2 Hz, 1H), 5.14 (dd, *J* = 7.4, 4.2 Hz, 1H), 3.53 (dt, *J* = 13.8, 5.0 Hz, 1H), 3.16 (dt, *J* = 13.8, 6.7 Hz, 1H), 2.84 – 2.79 (m, 1H), 1.98 (tt, *J* = 7.4, 2.4 Hz, 2H), 1.91 (ddt, *J* = 8.8, 7.2, 2.2 Hz, 2H), 1.60 – 1.51 (m, 4H). $^{13}$C NMR (151 MHz, DMSO-*d*$_6$) δ 170.9, 154.3, 130.9, 130.5, 130.4, 127.3, 121.7, 83.2, 78.9, 71.7, 44.3, 31.4, 29.3, 28.2, 28.2, 12.6. ESI$^+$ m/z found 284.15, calc'd 284.15 ($MH^+$-$N_2$).

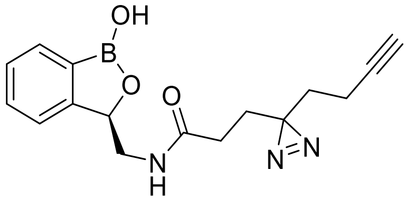

**(*R*)-3-(3-(But-3-yn-1-yl)-3*H*-diazirin-3-yl)-*N*-((1-hydroxy-1,3-dihydrobenzo[*c*][1,2]oxaborol-3-yl)methyl)propanamide ((*R*)-DMP1).**

An amber 1-dram vial under Ar was charged with 3-(3-(but-3-yn-1-yl)-3*H*-diazirin-3-yl)propanoic acid (92 mg, 0.55 mmol, 1.1 equiv), (*R*)-3-(aminomethyl)-benzo[*c*][1,2]oxaborol-1(3*H*)-ol hydrochloride (99.8 mg, 0.501 mmol), and HATU (285 mg, 0.750 mmol, 1.5 equiv). The mixture was cooled to 0 °C in an ice-water bath. Dry DMF (2.0 ml) and DIPEA (0.26 ml, 0.19 g, 1.5 mmol, 3.0 equiv) were added in that order with stirring. After 30 min, the cooling bath was removed, and the mixture was stirred at ambient temperature overnight. The mixture was diluted with EA (15 ml) and washed twice with $HCl_{(aq)}$ (1.0 M) and once with brine (15 ml each). The aqueous layers were extracted twice further with EA (20 ml each) in the same sequence. The organic layers were dried over $Na_2SO_4$ and filtered, and volatiles were removed at reduced pressure. The crude product was purified by flash column chromatography (isocratic 10% MeOH in DCM, then repurified with EA gradient in Hex, $R_f$ = 0.21 in 50% EA/Hex) followed by trituration with 1:2 ether/Hex (3 ml) at 4 °C. The solid was collected by vacuum filtration and washed twice with cold hexanes (1 ml each) to give the title compound as a white powder (61.9 mg, 0.310 mmol, 40% yield). NMR and ESI$^+$ spectra were consistent with those of the opposite enantiomer.

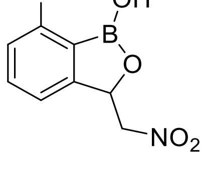

**7-Fluoro-3-(nitromethyl)benzo[*c*][1,2]oxaborol-1(3*H*)-ol.**

In a 100 ml round-bottom flask, 2-fluoro-6-formylphenylboronic acid (3.36 g, 20.1 mmol) was suspended in nitromethane (11 ml, 12 g, 200 mmol, 10 equiv). The mixture was cooled to 0 °C in an ice bath. Ice-cold $NaOH_{(aq)}$ (840 mg, 21.0 mmol, 1.05 equiv in 18 ml $H_2O$) was added slowly with stirring. The mixture was stirred at 0 °C for 1 h. Then, the cooling bath was removed, and the mixture was stirred at ambient temperature until TLC analysis (100%

EA) indicated high conversion of the starting aldehyde (3 hours). The mixture was cooled to 0 °C and quenched by adding $HCl_{(aq)}$ (3 M, 10 ml). The mixture was transferred to a separatory funnel and diluted with EA and $H_2O$ until each layer was approximately 50 ml. The layers were mixed and separated. The organic layer was washed with brine (50 ml). The aqueous layers were extracted further with EA (50 ml) in the same sequence. The organic layers were dried over $Na_2SO_4$ and filtered through cotton. Volatiles were removed at reduced pressure to give the crude product as a yellow solid (5.5 g). The crude product was purified by flash column chromatography (EA/Hex gradient; $R_f$ irreproducible on TLC, generally >0.5 in 100% EA) to give the title compound as a white solid (3.21 g, 15.2 mmol, 76% yield). $^1$H NMR (400 MHz, DMSO-$d_6$) δ 9.56 (s, 1H), 7.59 (td, $J$ = 7.9, 5.3 Hz, 1H), 7.39 (d, $J$ = 7.5 Hz, 1H), 7.13 (t, $J$ = 8.2 Hz, 1H), 5.81 (dd, $J$ = 9.0, 2.8 Hz, 1H), 5.34 (dd, $J$ = 13.7, 2.8 Hz, 1H), 4.69 (dd, $J$ = 13.6, 9.0 Hz, 1H). $^{19}$F NMR (376 MHz, DMSO) δ -104.8.

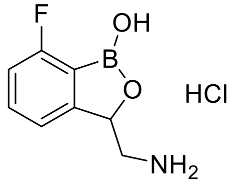

**(*rac*)-3-(Aminomethyl)-7-fluorobenzo[*c*][1,2]oxaborol-1(3*H*)-ol hydrochloride.**

In a 250 ml round-bottom flask under Ar, Raney Nickel (4 ml, 50% aqueous slurry, approximately 34 mmol, 2.2 equiv) was suspended in MeOH (15 ml) and cooled to 0 °C in an ice bath. Methanolic ammonia (7 M, 11 ml, 77 mmol, 5.1 equiv) was added with stirring. In a separate flask, 7-fluoro-3-(nitromethyl)benzo[*c*][1,2]oxaborol-1(3*H*)-ol (3.21 g, 15.2 mmol) was dissolved in MeOH (10 ml) and added to the reaction flask with stirring (final [$NH_3$] = 2 M). The atmosphere was purged with hydrogen, the cooling bath was removed, and the mixture was stirred at ambient temperature under an atmosphere of hydrogen (balloon pressure) overnight. The atmosphere was purged with Ar. The solution was filtered through a pad of Celite, which was rinsed thrice with MeOH (10 ml each). Volatiles were removed at reduced pressure. The residue was suspended in dry dioxane (30 ml). HCl (4 M in dioxane, 15 ml, 60 mmol, 4 equiv) was added with stirring. The cooling bath was removed, and the mixture was stirred at ambient temperature for 1 h. The mixture was cooled to 0 °C again, and the solid precipitate was collected by filtration and washed thrice with ether (10 ml each). The solid was dried under vacuum to give the crude racemate of the title compound as a light green powder (4.15 g, high mass due to Ni contamination).

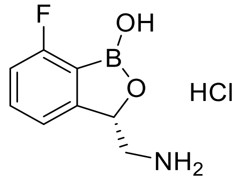

**(*S*)-3-(Aminomethyl)-7-fluorobenzo[*c*][1,2]oxaborol-1(3*H*)-ol hydrochloride.**

The crude racemic 3-(aminomethyl)-7-fluorobenzo[*c*][1,2]oxaborol-1(3*H*)-ol hydrochloride (4.15 g) obtained as described above was suspended in absolute EtOH (30 ml). Sodium hydroxide (610 mg, 15.3 mmol, 1.0 equiv) was added with stirring, and the mixture was stirred at ambient temperature for 1 h. Dry MeOH (30 ml) was added, and the mixture was filtered through a pad of Celite, which was rinsed thrice with 1:1 MeOH/EtOH (4 ml each), to give a clear blue solution. (*S*)-Mandelic acid ((*S*)-MA) (2.32 g, 15.3 mmol, 1.0 equiv) was added with stirring, and the mixture was stirred at ambient temperature for 20 min. The mixture was concentrated to approximately 15 ml at reduced pressure and then diluted again with absolute EtOH (20 ml), which caused a white solid to precipitate. This solid was collected by filtration, rinsed thrice with EtOH (4 ml each), and dried under vacuum to give the (*S*)-MA salt of title compound as a white powder (2.65 g, 7.96 mmol). Marfey analysis determined that this sample contained the (*S*) enantiomer in 68% ee. The filtrate was saved (see below).

This sample was recrystallized by dissolving in boiling 1:1 MeOH/EtOH (220 ml), filtering through a pad of Celite, and letting cool. The crystals were collected by filtration and rinsed twice with EtOH (10 ml each). The liquor was concentrated, and two further crops were collected in the same manner using lower volumes (100 ml, 60 ml). The three crops were combined to give the (*S*)-MA salt of the title compound as fluffy white solid (1.79 g, 5.37 mmol, 96% ee by Marfey analysis). This sample was dissolved in dioxane (12 ml). A dioxane solution of HCl (4 M, 3 ml) was added with stirring. The mixture was stirred at ambient temperature for 90 min to obtain a gel-like slurry. Ether (15 ml) and water (10 drops) were added with stirring; after approximately 30 min, a white precipitate formed. The solid was collected by filtration and rinsed thrice with ether (10 ml each) to give the title compound as a white powder (1.10 g, 5.06 mmol, 33% yield). $^{1}$H NMR (600 MHz, DMSO-$d_6$) δ 9.52 (s, 1H), 8.26 (s, 3H), 7.59 (td, *J* = 7.8, 5.2 Hz, 1H), 7.38 (d, *J* = 7.5 Hz, 1H), 7.14 (t, *J* = 8.1 Hz, 1H), 5.41 (dd, *J* = 8.6, 2.9 Hz, 1H), 3.48 (d, *J* = 13.2 Hz, 1H), 2.89 (t, *J* = 11.0 Hz, 1H). $^{13}$C NMR (151 MHz, DMSO-$d_6$) δ 163.4 (d, *J* = 250.4 Hz), 155.2 (d, *J* = 8.8 Hz), 134.2 (d, *J* = 7.2 Hz), 118.5 (d, *J* = 3.4 Hz), 117.6, 114.4 (d, *J* = 22.6 Hz), 76.8, 43.3. $^{19}$F NMR (376 MHz, DMSO) δ -104.5. ESI$^+$ m/z found 164.04, calc'd 164.07 ($MH^+$-$H_2O$). $[\alpha]_D^{23}$ = +51.2° (c 1.2, $H_2O$), compare to $[\alpha]_D^{31}$ = +71° (c 2.0, $H_2O$) reported for (*S*)-3-(aminomethyl)benzo[*c*][1,2]oxaborol-1(3*H*)-ol hydrochloride.[8, 9]

See above for procedure for Marfey analysis.

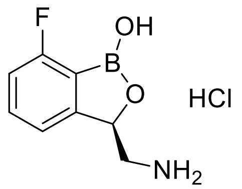

**(*R*)-3-(Aminomethyl)-7-fluorobenzo[*c*][1,2]oxaborol-1(3*H*)-ol hydrochloride.**

The saved filtrate from the first step of the previous compound was concentrated then redissolved in dioxane (15 ml). A dioxane solution of HCl (4 M, 7.5 ml) was added with stirring. The mixture was stirred at ambient temperature for 1 h to obtain a gel-like slurry. Ether (20 ml) was added with stirring; after approximately 30 min, a precipitate formed. The solid was collected by filtration and rinsed thrice with ether (15 ml each) to obtain a light green solid (1.96 g, 9.02 mmol). Marfey analysis determined that this sample contained the (*R*) enantiomer in 76% ee.

This sample was suspended in EtOH (20 ml) and basified by adding sodium hydroxide (310 mg, 7.75 mmol). The mixture was stirred at ambient temperature for 1 h, then diluted with MeOH (20 ml) and filtered through a plug of Celite, which was rinsed thrice with 1:1 MeOH/EtOH (4 ml each). (*R*)-MA (1.16 g, 7.62 mmol) was added with stirring, and the solution was allowed to sit at ambient temperature for 20 min. The solution was concentrated to approximately half the volume, which caused a precipitate to form. The solid was collected by filtration to obtain the (*R*)-MA salt of the title compound as a fluffy white solid (1.69 g, 5.07 mmol). The opposite enantiomer could not be detected by Marfey analysis, and this batch was therefore assigned as >98% ee. This sample was dissolved in dioxane (12 ml). A dioxane solution of HCl (4 M, 3 ml) was added with stirring. The mixture was stirred at ambient temperature for 90 min to obtain a gel-like slurry. Ether (15 ml) and water (10 drops) were added with stirring; after approximately 30 min, a white precipitate formed. The solid was collected by filtration and rinsed thrice with ether (10 ml each) to give the title compound as a white powder (1.08 g, 4.97 mmol, 33% yield). NMR and ESI$^+$ spectra were consistent with those of the opposite enantiomer. $[\alpha]_D^{23}$ = –45.6° (c 1.1, $H_2O$).

See above for procedure for Marfey analysis.

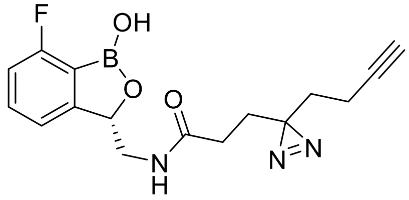

**(*S*)-3-(3-(But-3-yn-1-yl)-3*H*-diazirin-3-yl)-*N*-((7-fluoro-1-hydroxy-1,3-dihydrobenzo[*c*][1,2]oxaborol-3-yl)methyl)propanamide ((*S*)-DMP2).**

An amber 20 ml scintillation vial under Ar was charged with 3-(3-(but-3-yn-1-yl)-3*H*-diazirin-3-yl)propanoic acid (96 mg, 0.58 mmol, 1.2 equiv), (*S*)-3-(aminomethyl)-7-fluorobenzo[*c*][1,2]oxaborol-1(3*H*)-ol hydrochloride (109 mg, 0.501 mmol), and HATU (229 mg, 0.600 mmol, 1.2 equiv). The mixture was cooled to 0 °C in an ice-water bath. Dry DMF (2.0 ml) and DIPEA (0.26 ml, 0.19 g, 1.5 mmol, 3.0 equiv) were added in that order with stirring. After 30 min, the cooling bath was removed, and the mixture was stirred at ambient temperature overnight. The mixture was diluted with EA (20 ml) and washed with $HCl_{(aq)}$ (1.0 M) and brine (20 ml each). The aqueous layers were extracted twice further with EA (20 ml each) in the same sequence. The organic layers were dried over $Na_2SO_4$ and filtered, and volatiles were removed at reduced pressure. The crude product was purified by reverse-phase flash chromatography (C18, 20–90% MeCN gradient in $H_2O$ with 0.1% TFA, product-containing fractions identified by $KMnO_4$ stain and LC/MS analysis). Pure product-containing fractions were combined, concentrated, frozen, and lyophilized to give the title compound as a white powder (150 mg, 0.456 mmol, 91% yield). $^1$H NMR (600 MHz, DMSO-$d_6$) δ 9.28 (s, 1H), 8.03 (t, *J* = 5.7 Hz, 1H), 7.52 (td, *J* = 7.8, 5.2 Hz, 1H), 7.24 (d, *J* = 7.4 Hz, 1H), 7.06 (t, *J* = 8.1 Hz, 1H), 5.19 (dd, *J* = 6.7, 4.2 Hz, 1H), 3.50 (ddd, *J* = 13.9, 5.4, 4.2 Hz, 1H), 3.30 (dt, *J* = 13.9, 6.4 Hz, 1H), 2.81 (t, *J* = 2.6 Hz, 1H), 1.97 (td, *J* = 7.4, 2.7 Hz, 2H), 1.94 – 1.84 (m, 2H), 1.61 – 1.50 (m, 4H). $^{13}$C NMR (151 MHz, DMSO-$d_6$) δ 171.0, 163.3 (d, *J* = 249.9 Hz), 157.2 (d, *J* = 8.5 Hz), 133.7 (d, *J* = 7.2 Hz), 118.3 (d, *J* = 3.3 Hz), 117.8, 113.7 (d, *J* = 22.6 Hz), 83.1, 78.71, 71.7, 43.8, 31.4, 29.3, 28.2, 28.1, 12.6. $^{19}$F NMR (376 MHz, DMSO) δ -105.2. ESI$^+$ m/z found 302.15, calc'd 302.14 ($MH^+$-$N_2$).

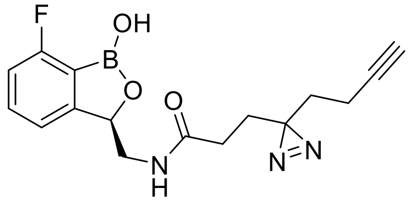

**(*R*)-3-(3-(But-3-yn-1-yl)-3*H*-diazirin-3-yl)-*N*-((7-fluoro-1-hydroxy-1,3-dihydrobenzo[*c*][1,2]oxaborol-3-yl)methyl)propanamide ((*R*)-DMP2).**

An amber 20 ml scintillation vial under Ar was charged with 3-(3-(but-3-yn-1-yl)-3*H*-diazirin-3-yl)propanoic acid (96 mg, 0.58 mmol, 1.2 equiv), (*R*)-3-(aminomethyl)-7-fluorobenzo[*c*][1,2]oxaborol-1(3*H*)-ol hydrochloride (109 mg, 0.501 mmol), and HATU (229 mg, 0.600 mmol, 1.2 equiv). The mixture was cooled to 0 °C in an ice-water bath. Dry DMF (2.0 ml) and DIPEA (0.26 ml, 0.19 g, 1.5 mmol, 3.0 equiv) were added in that order with stirring. After 30 min, the cooling bath was removed, and the mixture was stirred at ambient temperature overnight. The mixture was diluted with EA (20 ml) and washed with $HCl_{(aq)}$ (1.0 M) and brine (20 ml each). The aqueous layers were extracted twice further with EA (20 ml each) in the same sequence. The organic layers were dried over $Na_2SO_4$ and filtered, and volatiles were removed at reduced pressure. The crude product was purified by reverse-phase flash chromatography (C18, 20–90% MeCN gradient in $H_2O$ with 0.1% TFA, product-containing fractions identified by $KMnO_4$ stain and LC/MS analysis). Pure product-containing fractions were combined, concentrated, frozen, and lyophilized to give the title compound as a white powder (131 mg, 0.398 mmol, 79% yield). NMR and ESI$^+$ spectra were consistent with those of the opposite enantiomer.

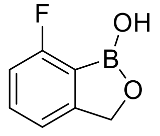

**7-Fluorobenzo[*c*][1,2]oxaborol-1(3*H*)-ol.**

In a 1-dram vial, (2-fluoro-6-formyl-phenyl)boronic acid (168 mg, 1.00 mmol) was dissolved in dry MeOH (2.0 ml). The solution was cooled to 0 °C in an ice-water bath. Sodium borohydride (45 mg, 1.2 mmol, 1.2 equiv) was added with stirring, and the mixture was stirred at 0 °C for 1 h. Then the cooling bath was removed, and the mixture was stirred at ambient temperature overnight. LC/MS analysis indicated incomplete conversion. The mixture was cooled to 0 °C and an additional portion of sodium borohydride (23 mg, 0.61 mmol, 0.61 equiv) was added with stirring. The cooling bath was removed, and the mixture was stirred at ambient temperature for an additional 1 h. The reaction was quenched with $HCl_{(aq)}$ (3 M, 0.1 ml). The mixture was concentrated then diluted with water (1.5 ml). The mixture was extracted thrice with EA (2 ml each). The organic layers were dried over $Na_2SO_4$ and filtered, and volatiles were removed at reduced pressure. The crude product was purified by flash column chromatography (0–60% EA gradient in Hex) to give the title compound as a white solid (131 mg, 0.862 mmol, 86% yield). $^1$H NMR (400 MHz, DMSO-$d_6$) δ 9.26 (s, 1H), 7.53 (td, $J$ = 7.8, 5.3 Hz, 1H), 7.24 (d, $J$ = 7.5 Hz, 1H), 7.05 (t, $J$ = 8.2 Hz, 1H), 5.01 (s, 2H). $^{19}$F NMR (376 MHz, DMSO) δ -105.1.

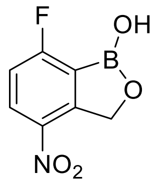

**7-Fluoro-4-nitrobenzo[*c*][1,2]oxaborol-1(3*H*)-ol.**

In a 10 ml round bottom flask, fuming nitric acid was cooled to –40 °C in a MeCN/dry-ice bath. 7-Fluorobenzo[*c*][1,2]oxaborol-1(3*H*)-ol (277 mg, 1.82 mmol) was added slowly with stirring over the course of 10 min. The mixture was stirred at –40 °C for 1 h. The mixture was poured onto ice, then diluted with water (2 ml) to obtain a precipitate. The solid was collected by vacuum filtration, rinsed with cold water (2 ml), and dried under vacuum to obtain the crude product (171 mg). $^1$H NMR analysis of this sample indicated a 10:1 ratio of isomers. The crude product was recrystallized by dissolving in hot dilute $HCl_{(aq)}$ (1 drop 6 M $HCl_{(aq)}$ in 30 ml water), filtering through a pad of Celite, allowing the solution to slowly cool, then storing at 4 °C overnight. The solid was collected by vacuum filtration, rinsed thrice with cold water (2 ml each), and dried under vacuum to give the title compound as a white powder (85.0 mg, 0.432 mmol, 24% yield, >100:1 isomer ratio). $^1$H NMR (400 MHz, DMSO-$d_6$) δ 9.65 (s, 1H), 8.42 (dd, $J$ = 8.9, 4.3 Hz, 1H), 7.41 (dd, $J$ = 8.9, 7.1 Hz, 1H), 5.38 (s, 2H). $^{19}$F NMR (376 MHz, DMSO) δ -94.7.

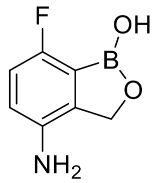

**4-Amino-7-fluorobenzo[*c*][1,2]oxaborol-1(3*H*)-ol.**

In a 1-dram vial under Ar, 7-fluoro-4-nitrobenzo[*c*][1,2]oxaborol-1(3*H*)-ol (82.7 mg, 0.420 mmol) was dissolved in IPA (0.8 ml) and cooled to 0 °C in an ice-water bath. In a separate 1-dram vial under Ar, Pd/C (10% wt, 45 mg, 0.042 mmol, 0.10 equiv) was suspended in IPA and cooled to 0 °C in an ice-water bath. Aqueous ammonium formate (265 mg, 4.20 mmol, 10 equiv, in 0.4 ml water) was added to the Pd/C suspension with stirring, then the resulting suspension was added to the reaction vial with stirring. The mixture was stirred at 0 °C for 1 h. The mixture was filtered through a pad of Celite, which was rinsed thrice with IPA. The mixture was concentrated then partitioned between EA and water (5 ml each). The layers were separated, and the organic layer was washed again with brine (5 ml). The aqueous layers were extracted twice further in the same sequence with EA (5 ml each). The organic layers were dried over $Na_2SO4$ and filtered. Volatiles were removed at reduced pressure to give the title compound as an off-white solid (71.6 mg, 0.429 mmol, quant yield). $^1$H NMR (400 MHz, Methanol-$d_4$) δ 6.84 – 6.69 (m, 2H), 4.94 (s, 2H). $^{19}$F NMR (376 MHz, MeOD) δ -120.8. ESI$^+$ m/z found 168.08, calc'd 168.06 ($MH^+$).

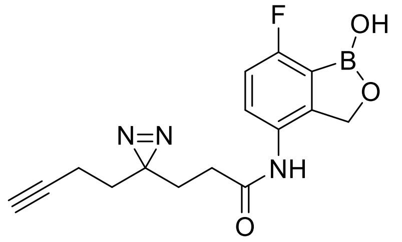

**3-(3-(But-3-yn-1-yl)-3*H*-diazirin-3-yl)-*N*-(7-fluoro-1-hydroxy-1,3-dihydrobenzo[*c*][1,2]oxaborol-4-yl)propanamide (DMP3).**

An amber 1-dram vial under Ar was charged with 3-(3-(but-3-yn-1-yl)-3*H*-diazirin-3-yl)propanoic acid (56.3 mg, 0.339 mmol, 1.1 equiv), 4-amino-7-fluorobenzo[*c*][1,2]oxaborol-1(3*H*)-ol (50.0 mg, 0.299 mmol), and HATU (171 mg, 0.450 mmol, 1.5 equiv). The mixture was cooled to 0 °C in an ice-water bath. Dry DMF (1.2 ml) and DIPEA (0.16 ml, 0.12 g, 1.5 mmol, 3.1 equiv) were added in that order with stirring. After 30 min, the cooling bath was removed, and the mixture was stirred at ambient temperature overnight. The mixture was diluted with EA (5 ml) and washed twice with $HCl_{(aq)}$ (1.0 M) and once with brine (5 ml each). The aqueous layers were extracted twice further with EA (5 ml each) in the same sequence. The organic layers were dried over $Na_2SO_4$ and filtered, and volatiles were removed at reduced pressure. The crude product was redissolved in 1:1 MeCN/$H_2O$ with 0.1% v/v FA (3 ml) and filtered (0.25 μm PTFE), and the filter was rinsed with an additional portion of the same solvent (3 ml). The resulting solution was purified by preparative reverse-phase HPLC (C18, 5–95% MeCN gradient in $H_2O$ with 0.1% FA, product-containing fractions identified with a split-line UV/vis detector). Pure product-containing fractions were combined, and volatiles were removed in a centrifugal evaporator (Genevac) to give the title compound as a white powder (50.8 mg, 0.161 mmol, 54% yield). $^1$H NMR (600 MHz, DMSO-$d_6$) δ 9.55 (s, 1H), 9.26 (s, 1H), 7.59 (dd, *J* = 8.6, 4.5 Hz, 1H), 7.05 (t, *J* = 8.0 Hz, 1H), 4.94 (s, 2H), 2.83 (t, *J* = 2.6 Hz, 1H), 2.15 (t, *J* = 7.6 Hz, 2H), 2.02 (td, *J* = 7.4, 2.7 Hz, 2H), 1.75 (t, *J* = 8.2, 7.7 Hz, 2H), 1.62 (t, *J* = 7.4 Hz, 2H). $^{13}$C NMR (151 MHz, DMSO-$d_6$) δ 169.6, 160.4 (d, *J* = 247.1 Hz), 148.6 (d, *J* = 8.9 Hz), 128.4 (d, *J* = 3.2 Hz), 127.9 (d, *J* = 7.7 Hz), 117.8, 113.8 (d, *J* = 24.3 Hz), 83.2, 71.8, 69.1, 31.5, 29.8, 28.2, 27.8, 12.7. $^{19}$F NMR (376 MHz, DMSO) δ -110.28. ESI$^+$ m/z found 288.13, calc'd 288.12 ($MH^+$-$N_2$).

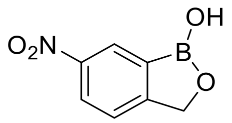

**6-Nitrobenzo[*c*][1,2]oxaborol-1(3*H*)-ol.**

In a 50 ml round bottom flask, fuming nitric acid was cooled to –40 °C in a MeCN/dry-ice bath. Benzo[*c*][1,2]oxaborol-1(3*H*)-ol (1.34 g, 10.0 mmol) was added in small portions with stirring over the course of 1 h to obtain a light yellow suspension. The mixture was poured into cold water (50 ml) in an ice-water bath. After mixing well, the solid was collected by vacuum filtration and rinsed thrice with cold water (30 ml each). The resulting wet solid was frozen and lyophilized to give the title compound as a free-flowing white powder (1.59 g, 8.88 mmol, 89% yield). $^1$H NMR (400 Hz, DMSO-$d_6$) δ 9.58 (s, 1H), 8.59 (d, *J* = 2.3 Hz, 1H), 8.33 (dd, *J* = 8.4, 2.3 Hz, 1H), 7.70 (d, *J* = 8.4 Hz, 1H), 5.13 (s, 2H).

Note: an impurity observed by $^1$H NMR spectroscopy likely corresponds to a minor isomer of the product (1:43 ratio).

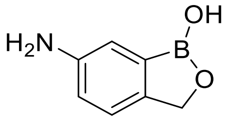

**6-Aminobenzo[*c*][1,2]oxaborol-1(3*H*)-ol.**

In a 50 ml round bottom flask under Ar, Pd/C (10% wt, 532 mg, 0.500 mmol, 0.10 equiv) was suspended in IPA. The mixture was cooled to 0 °C in an ice-water bath. Aqueous ammonium formate (3.16 g, 50.1 mmol, 10 equiv in 5.0 ml water) was added with stirring. 6-Nitrobenzo[*c*][1,2]oxaborol-1(3*H*)-ol (895 mg, 5.00 mmol) was added with stirring. The mixture was stirred at 0 °C for 3 h. The mixture was filtered through a pad of Celite then concentrated to approximately 5 ml. The mixture was partitioned between EA and water (20 ml each). The layers were mixed and separated. The organic layer was washed with brine (20 ml). The aqueous layers were extracted twice further in the same sequence with EA (20 ml each). The organic layers were dried over $Na_2SO_4$ and filtered. The mixture was concentrated at reduced pressure. The sample was dissolved in MeOH, then concentrated again. The sample was triturated with wet ether (10 ml, required 3 cycles of adding either and evaporating to obtain suitable solid). The solid was collected by vacuum filtration and rinsed twice with cold ether (4 ml each) to give the title compound as an off-white solid (547 mg, 3.67 mmol, 73% yield). $^1$H NMR (400 MHz, DMSO-$d_6$) δ 8.89 (s, 1H), 7.03 (d, *J* = 8.1 Hz, 1H), 6.88 (d, *J* = 2.2 Hz, 1H), 6.70 (dd, *J* = 8.0, 2.2 Hz, 1H), 4.97 (s, 2H), 4.81 (s, 2H).

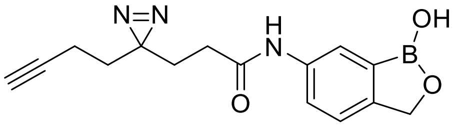

**3-(3-(But-3-yn-1-yl)-3*H*-diazirin-3-yl)-*N*-(1-hydroxy-1,3-dihydrobenzo[*c*][1,2]oxaborol-6-yl)propanamide (DMP4).**

In an amber 1-dram vial under Ar, 3-(3-(but-3-yn-1-yl)-3*H*-diazirin-3-yl)propanoic acid (91.9 mg, 0.550 mmol, 1.1 equiv) was dissolved in dry DMF (2.0 ml). The solution was cooled to 0 °C in an ice-water bath. 6-Aminobenzo[*c*][1,2]oxaborol-1(3*H*)-ol (74.5 mg, 0.500 mmol), HATU (285 mg, 0.750 mmol, 1.5 equiv), and DIPEA (0.13 ml, 96 mg, 0.75 mmol, 1.5 equiv) were added in that order with stirring. After 30 min, the cooling bath was removed, and the mixture was stirred at ambient temperature overnight. The mixture was diluted with EA (8 ml) and washed with $HCl_{(aq)}$ (1.0 M) and brine (8 ml each). The aqueous layers were extracted twice further with EA (8 ml each) in the same sequence. The organic layers were dried over $Na_2SO_4$ and filtered, and volatiles were removed at reduced pressure. The crude product was purified by column chromatography (hand-packed column with ~100 ml silica slurry, 10% MeOH/DCM isocratic, Rf 0.51 in 10% MeOH/DCM) to give the title compound as a light yellow solid (97.2 mg, 0.327 mmol, 65% yield). $^1$H NMR (600 MHz, DMSO-$d_6$) δ 9.92 (s, 1H), 9.18 (s, 1H), 7.98 (d, *J* = 2.0 Hz, 1H), 7.58 (dd, *J* = 8.2, 2.0 Hz, 1H), 7.32 (d, *J* = 8.2 Hz, 1H), 4.93 (s, 2H), 2.87 – 2.78 (m, 1H), 2.15 (t, *J* = 7.3 Hz, 2H), 2.02 (td, *J* = 7.4, 2.7 Hz, 2H), 1.76 (t, *J* = 7.9 Hz, 2H), 1.61 (t, *J* = 7.4 Hz, 2H). $^{13}$C NMR (151 MHz, DMSO) δ 169.6, 148.6, 137.9, 130.9, 122.2, 121.4, 121.0, 83.2, 71.8, 69.6, 31.5, 30.4, 28.3, 27.8, 12.7. ESI$^+$ m/z found 270.14, calc'd 270.13 ($MH^+$-$N_2$).

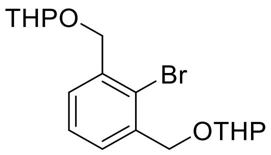

**2,2'-(((2-Bromo-1,3-phenylene)bis(methylene))bis(oxy))bis(tetrahydro-2*H*-pyran).**

In a 20 ml scintillation vial, 2-bromobenzene-1,3-dicarbaldehyde (426 mg, 2.00 mmol) was dissolved in dry MeOH (4.0 ml). Sodium borohydride (158 mg, 4.18 mmol, 2.1 equiv) was added slowly with stirring. The mixture was stirred at ambient temperature for 1 h. The reaction was quenched by slowly adding $HCl_{(aq)}$ (3 M, 1 ml). The mixture was diluted with water (30 ml) and extracted four times with ether (30 ml each). The organic layers were dried over $MgSO_4$ and filtered. Volatiles were removed at reduced pressure to give crude [2-bromo-3-(hydroxymethyl)phenyl]methanol. In a 100 ml round bottom flask, this sample was suspended in dry DCM (10 ml). Tosic acid monohydrate (38 mg, 0.20 mmol, 0.10 equiv) was added. Then, dihydropyran (0.55 ml, 0.51 g, 6.0 mmol, 3.0 equiv) was added slowly over the course of 30 min. The mixture was stirred at ambient temperature until a clear solution was obtained. The mixture was diluted with ether to approximately 20 ml and washed with $K_2CO_{3(aq)}$ (1 M) and brine (20 ml each). The aqueous layers were extracted in the same sequence with ether (20 ml). The organic layers were dried over $Na_2SO_4$ and filtered, and volatiles were removed at reduced pressure. The crude product was purified by flash column chromatography (EA gradient in Hex) to give the title compound as a clear and colorless oil (594 mg, 1.54 mmol, 77% yield). $^1$H NMR (400 MHz, Chloroform-*d*) δ 7.45 (d, *J* = 7.2 Hz, 2H), 7.33 (dd, *J* = 8.1, 7.0 Hz, 1H), 4.85 (d, *J* = 13.4 Hz, 2H), 4.79 (t, *J* = 3.5 Hz, 2H), 4.61 (d, *J* = 13.4 Hz, 2H), 3.93 (ddd, *J* = 11.4, 8.9, 3.2 Hz, 2H), 3.57 (dtd, *J* = 11.2, 4.4, 1.7 Hz, 2H), 1.98 – 1.83 (m, 2H), 1.83 – 1.55 (m, 10H).

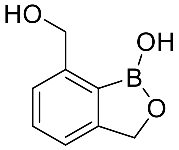

**7-(Hydroxymethyl)benzo[*c*][1,2]oxaborol-1(3*H*)-ol.**

In an oven-dried 10 ml round bottom flask under Ar, 2,2'-(((2-bromo-1,3-phenylene)bis(methylene))bis(oxy))bis(tetrahydro-2*H*-pyran) (538 mg, 1.40 mmol) was dissolved in dry THF (5.0 ml). The solution was cooled to –78 °C in an acetone/dry-ice bath. A solution of *n*-butyllithium in hexanes (2.5 M, 0.58 ml, 1.5 mmol, 1.0 equiv) was added slowly with stirring. The mixture was stirred at –78 °C for 30 min. Trimethylborate (0.47 ml, 0.44 g, 4.2 mmol, 3.0 equiv) was added slowly with stirring. After 10 min, the cooling bath was removed, and the mixture was stirred at ambient temperature for 3 h. The reaction was quenched by adding saturated $NaHCO_{3(aq)}$. The mixture was partitioned between EA and water (20 ml each). The layers were separated, and the organic layer was washed again with brine (20 ml). The aqueous layers were extracted again with EA (20 ml) in the same sequence. The organic layers were dried over $Na_2SO_4$ and filtered, and volatiles were removed at reduced pressure to give crude 2,6-bis(tetrahydropyran-2-yloxymethyl)phenyl]boronic acid. The sample was dissolved in MeOH (5.0 ml). Tosic acid monohydrate (13 mg, 0.068 mmol, 0.05 equiv) was added with stirring, and the mixture was stirred at ambient temperature for 2 h. The mixture was concentrated then partitioned between DCM and saturated $NaHCO_{3(aq)}$ (10 ml each). The layers were separated, and the organic layer was washed with brine. The aqueous layers were extracted further with DCM (10 ml) in the same sequence. The organic layers were filtered, and volatiles were removed at reduced pressure. The crude product was redissolved in EA (10 ml) and let sit to obtain a white precipitate. The solid was collected by vacuum filtration and rinsed twice with EA (2 ml each) to give the title compound as a white powder (119 mg, 0.726 mmol, 52% yield). $^1$H NMR (400 MHz, Methanol-$d_4$) δ 7.45 (t, *J* = 7.5 Hz, 1H), 7.42 – 7.22 (m, 2H), 5.18 – 4.94 (m, 2H), 4.93 – 4.70 (m, 2H).

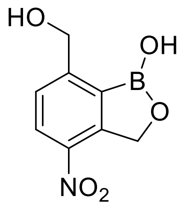

**7-(Hydroxymethyl)-4-nitrobenzo[*c*][1,2]oxaborol-1(3*H*)-ol.**

In a 1-dram vial, fuming nitric acid was cooled to –40 °C in a MeCN/dry-ice bath. 7-(Hydroxymethyl)benzo[*c*][1,2]oxaborol-1(3*H*)-ol (82.0 mg, 0.500 mmol) was added with stirring in small portions over the course of one hour. After the addition was complete, the mixture was stirred for an additional 1 h. The reaction was quenched by pouring the solution onto a few pieces of ice in a 20 ml vial, then the mixture was diluted further with water (1 ml). The mixture was extracted with EA (4 ml), and the organic layer was washed with brine (3 ml). The aqueous layers were extracted twice further in the same sequence with EA (4 ml each). The organic layers were dried by filtering through $Na_2SO4$, and volatiles were removed at reduced pressure. The crude product was loaded onto a plug of silica (ca. 2 ml) and eluted with 1:1 EA/Hex. Product-containing fractions were combined, concentrated, and dried under vacuum to give the title compound as an off-white solid (107.5 mg, 0.510 mmol, 103% yield). $^1$H NMR (400 MHz, DMSO-$d_6$) δ 9.63 (s, 1H), 8.37 (d, *J* = 8.3 Hz, 1H), 7.71 (d, *J* = 8.3 Hz, 1H), 5.86 (s, 2H), 5.42 (s, 2H).

Note: the aliphatic alcohol OH resonance was not observed in this $^1$H NMR spectrum; two interconverting isomers are possible for this structure of this compound.

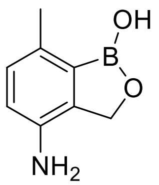

**4-Amino-7-methylbenzo[*c*][1,2]oxaborol-1(3*H*)-ol.**

In a 20 ml vial 7-(hydroxymethyl)-4-nitrobenzo[*c*][1,2]oxaborol-1(3*H*)-ol 105 mg, 0.500 mmol) was dissolved in IPA (1 ml), and the solution was cooled to 0 °C in an ice-water bath. In a separate 1-dram vial, palladium on carbon (10% wt, 53.5 mg, 0.0503 mmol, 0.10 equiv) was suspended in IPA (1 ml), and the mixture was cooled to 0 °C in an ice-water bath. In a separate 1-dram vial, ammonium formate (317 mg, 5.03 mmol, 10 equiv) was dissolved in water (0.5 ml). The aqueous ammonium formate solution was added to the palladium suspension with stirring, then the resulting mixture was added to the reaction vial with stirring. The vial was capped, and the mixture was stirred at 0 °C for 90 min, then the cooling bath was removed, and the mixture was stirred at ambient temperature overnight. The mixture was filtered through a pad of Celite, which was rinsed thrice with IPA (1 ml each). Volatiles were evaporated under a stream of Ar. The residue was partitioned between EA and water (5 ml each). The layers were separated, and the aqueous layer was extracted twice further with EA (5 ml each). The organic layers were concentrated at reduced pressure, and the crude product was purified by preparative RP-HPLC (C18, 5–95% MeCN gradient in $H_2O$ with 0.1% FA) to give the title compound as a white solid (39.3 mg, 0.240 mmol, 48% yield). $^1$H NMR (400 MHz, DMSO-$d_6$) δ 8.67 (s, 1H), 6.80 (d, *J* = 7.8 Hz, 1H), 6.55 (d, *J* = 7.8 Hz, 1H), 4.94 – 4.52 (m, 4H), 2.27 (s, 3H). ESI$^+$ m/z found 164.10, calc'd 164.09 ($MH^+$).

Note: the regioselectivity of this reaction was assigned based on the product of the next step.

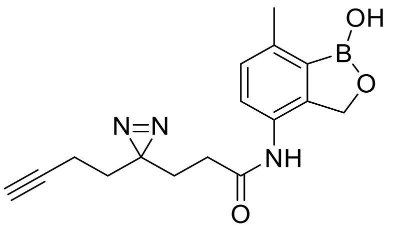

**3-(3-(But-3-yn-1-yl)-3*H*-diazirin-3-yl)-*N*-(1-hydroxy-7-methyl-1,3-dihydrobenzo[*c*][1,2]oxaborol-4-yl)propanamide (DMP5).**

In a 1-dram vial under Ar, 4-amino-7-methylbenzo[*c*][1,2]oxaborol-1(3*H*)-ol (29.5 mg, 0.180 mmol), HATU (93.9 mg, 0.250 mmol, 1.4 equiv), and 3-(3-(but-3-yn-1-yl)-3*H*-diazirin-3-yl)propanoic acid (32.9 mg, 0.200 mmol, 1.1 equiv) were cooled to 0 °C in an ice-water bath. DMF (0.66 ml) and DIPEA (8.6 µl, 0.49 mmol, 2.7 equiv) were added in that order with stirring. The mixture was stirred at 0 °C for 30 min, then at ambient temperature overnight. The mixture was partitioned between EA (4 ml) and $HCl_{(aq)}$ (1.0 M, 4 ml). The layers were separated, and the organic layer was washed further with $HCl_{(aq)}$ (1.0 M) and brine (4 ml each). The aqueous layers were extracted five times further in the same sequence with EA (4 ml each). Volatiles were removed at reduced pressure, and the crude product was purified by flash column chromatography (10% MeOH/DCM). Product containing fractions were concentrated, then the residue was triturated with wet ether (1 ml ether and 20 µl water) at 4 °C; the white precipitate was collected by filtration, washed twice further with cold ether (1 ml each). The sample was dried under vacuum to give the title compound as a white powder (38.7 mg, 0.124 mmol, 69% yield). $^1$H NMR (600 MHz, DMSO-$d_6$) δ 9.4 (s, 1H), 8.9 (s, 1H), 7.4 (d, *J* = 7.9 Hz, 1H), 7.1 (d, *J* = 7.9 Hz, 1H), 4.9 (s, 2H), 2.8 (t, *J* = 2.7 Hz, 1H), 2.4 (s, 3H), 2.1 (t, *J* = 7.6 Hz, 2H), 2.0 (td, *J* = 7.3, 2.6 Hz, 2H), 1.8 (t, *J* = 7.6 Hz, 2H), 1.6 (t, *J* = 7.4 Hz, 2H). $^{13}$C NMR (151 MHz, DMSO-$d_6$) δ 169.4, 146.1, 137.9, 130.6 – 130.2 (m), 129.5, 128.2, 125.2, 83.2, 71.7, 68.8, 31.5, 29.9, 28.3, 27.9, 19.3, 12.7. ESI$^+$ m/z found 284.15, calc'd 284.15 ($MH^+$-$N_2$).

Note: the structure of this product was assigned by 2D NMR spectroscopy (HSQC and HMBC); the methyl group was strongly correlated to one aromatic CH group.

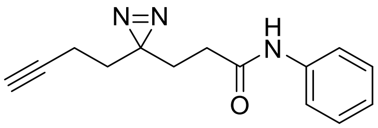

**3-(3-(But-3-yn-1-yl)-3*H*-diazirin-3-yl)-*N*-phenylpropanamide (C6).**

3-(3-(But-3-yn-1-yl)-3*H*-diazirin-3-yl)propanoic acid (46.6 mg, 0.500 mmol) was dissolved in dry DMF (2.0 ml) in an amber 1-dram vial, and the solution was cooled to 0 °C in an ice-water bath. Aniline (46 µl, 0.55 mmol, 1.1 equiv), HATU (285 mg, 0.750 mmol, 1.5 equiv), and DIPEA (0.26 ml, 1.5 mmol, 3.0 equiv) were added in that order with stirring. The mixture was stirred at 0 °C for approximately 30 min, then the cooling bath was removed, and the mixture was stirred at ambient temperature overnight. The mixture was diluted with ethyl acetate (8 ml) and washed with water, 1 M $HCl_{(aq)}$, and brine (8 ml each) in that order. The aqueous layers were extracted twice further with ethyl acetate (8 ml each) in the same sequence. The organic layers were filtered through $Na_2SO_4$, and volatiles were removed at reduced pressure. The crude product was purified by flash column chromatography (0–10% MeOH gradient in DCM, then repurified with 0–20% ethyl acetate gradient in Hex, Rf 0.30 in 20% EA/Hex) to give the title compound as a white solid (97.2 mg, 0.403 mmol, 81% yield). $^1$H NMR (400 MHz, $CDCl_3$) δ 7.49 (d, *J* = 7.9 Hz, 2H), 7.33 (t, *J* = 7.9 Hz, 2H), 7.12 (t, *J* = 7.4 Hz, 1H), 7.08 (s, 1H), 2.12 (t, *J* = 7.5 Hz, 2H), 2.05 (td, *J* = 7.4, 2.6 Hz, 2H), 1.99 (t, *J* = 2.6 Hz, 1H), 1.95 (t, *J* = 7.5 Hz, 2H), 1.69 (t, *J* = 7.4 Hz, 2H).

*Chemical abbreviations*

EA – ethyl acetate

Hex – hexanes (mixture of isomers)

THF – tetrahydrofuran

DCM – dichloromethane

DMF – dimethylformamide

DMSO – dimethylsulfoxide

IPA – *iso*-propanol

MeCN – acetonitrile

TFA – trifluoroacetic acid

FA – formic acid

DIPEA – di-*iso*-propylethylamine

HATU – 1-[bis(dimethylamino)methylene]-1H-1,2,3-triazolo[4,5-b]pyridinium 3-oxide hexafluorophosphate

MeOH – methanol

EtOH – ethanol

**(*S*)-DMP1.** $^{1}$H NMR (600 MHz, DMSO-$d_6$)

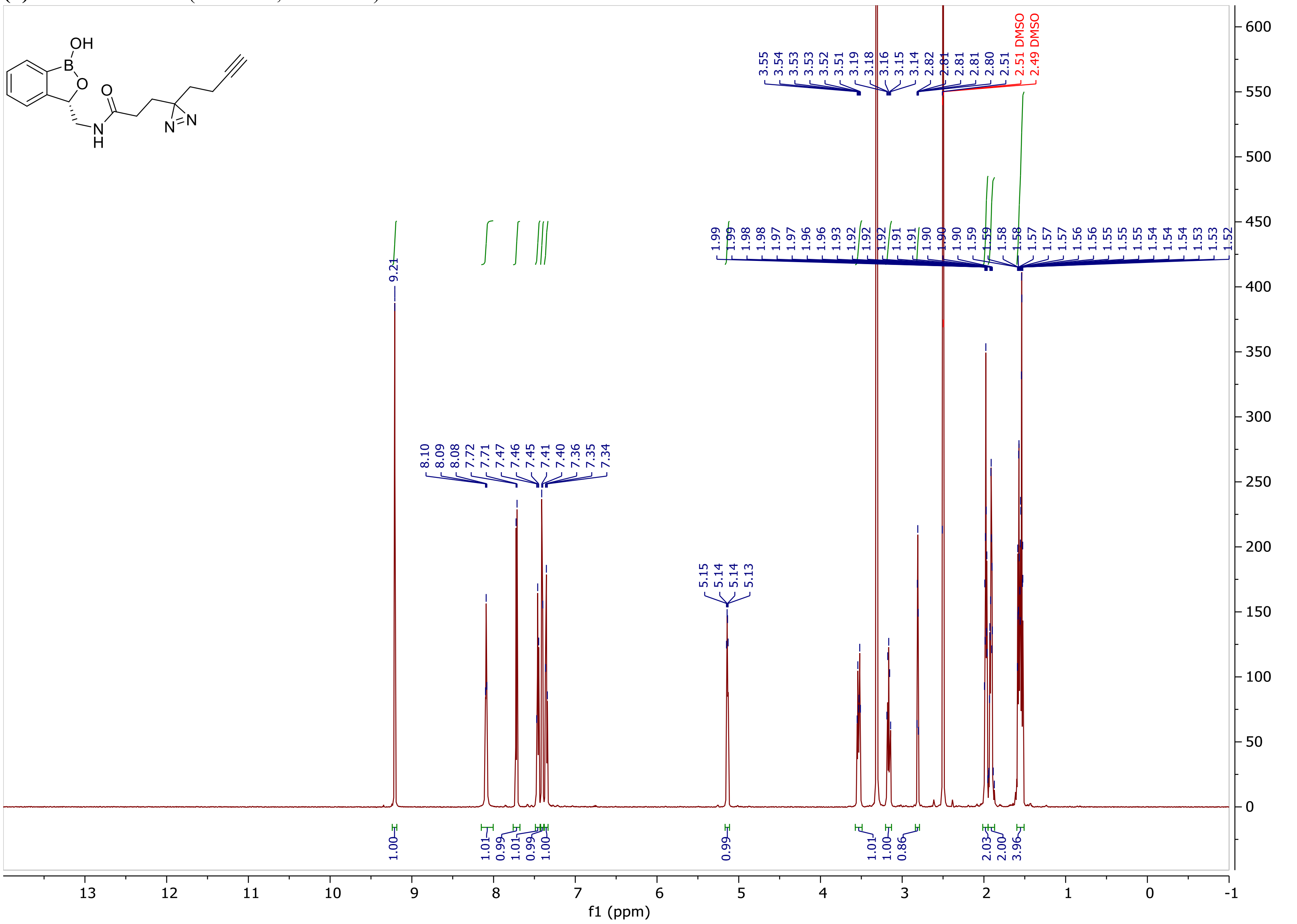

(***S***)**-DMP1.** $^{13}$C NMR (151 MHz, DMSO-$d_6$)

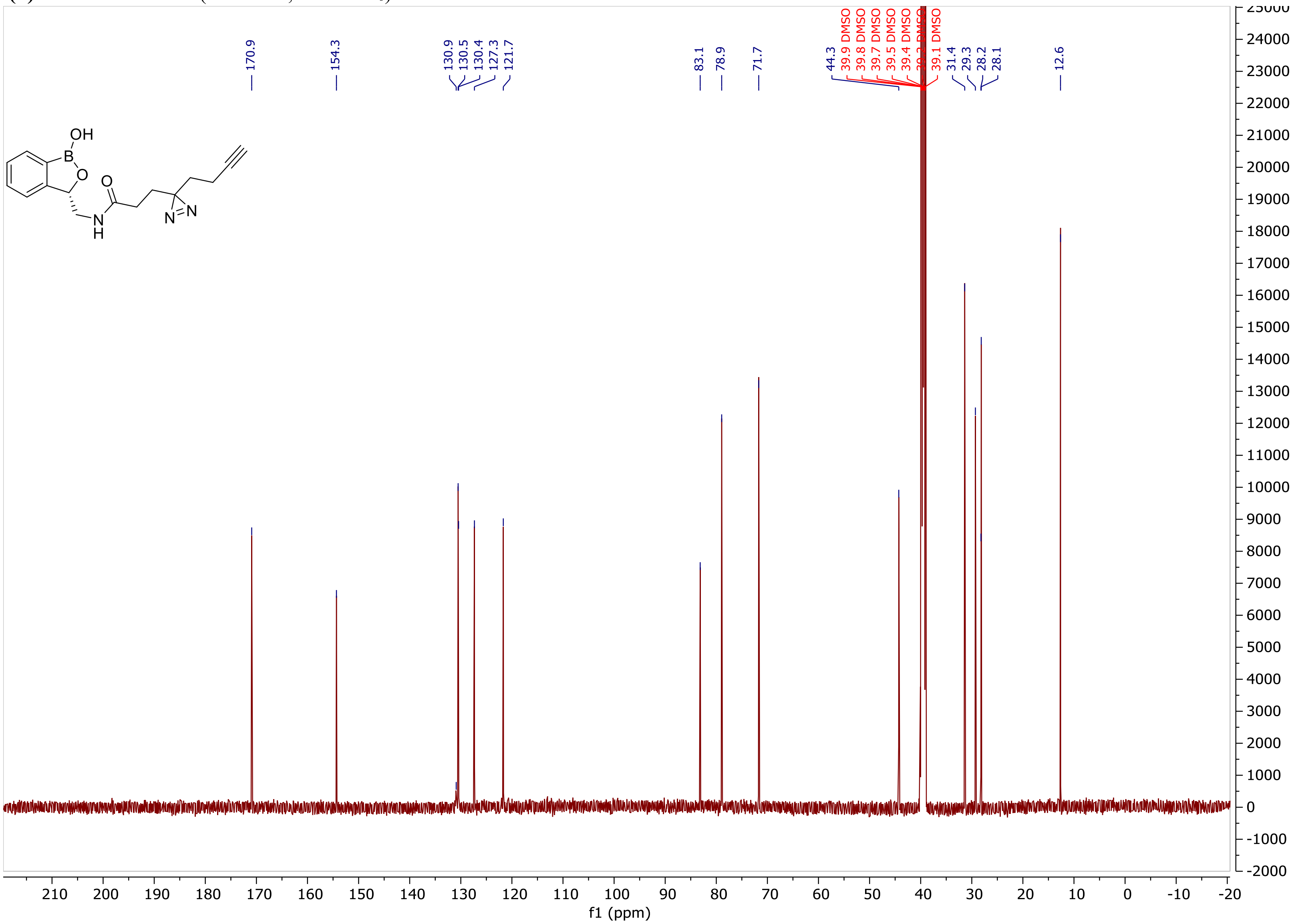

**(*S*)-3-(Aminomethyl)-7-fluorobenzo[*c*][1,2]oxaborol-1(3*H*)-ol hydrochloride.** $^1$H NMR (600 MHz, DMSO-$d_6$)

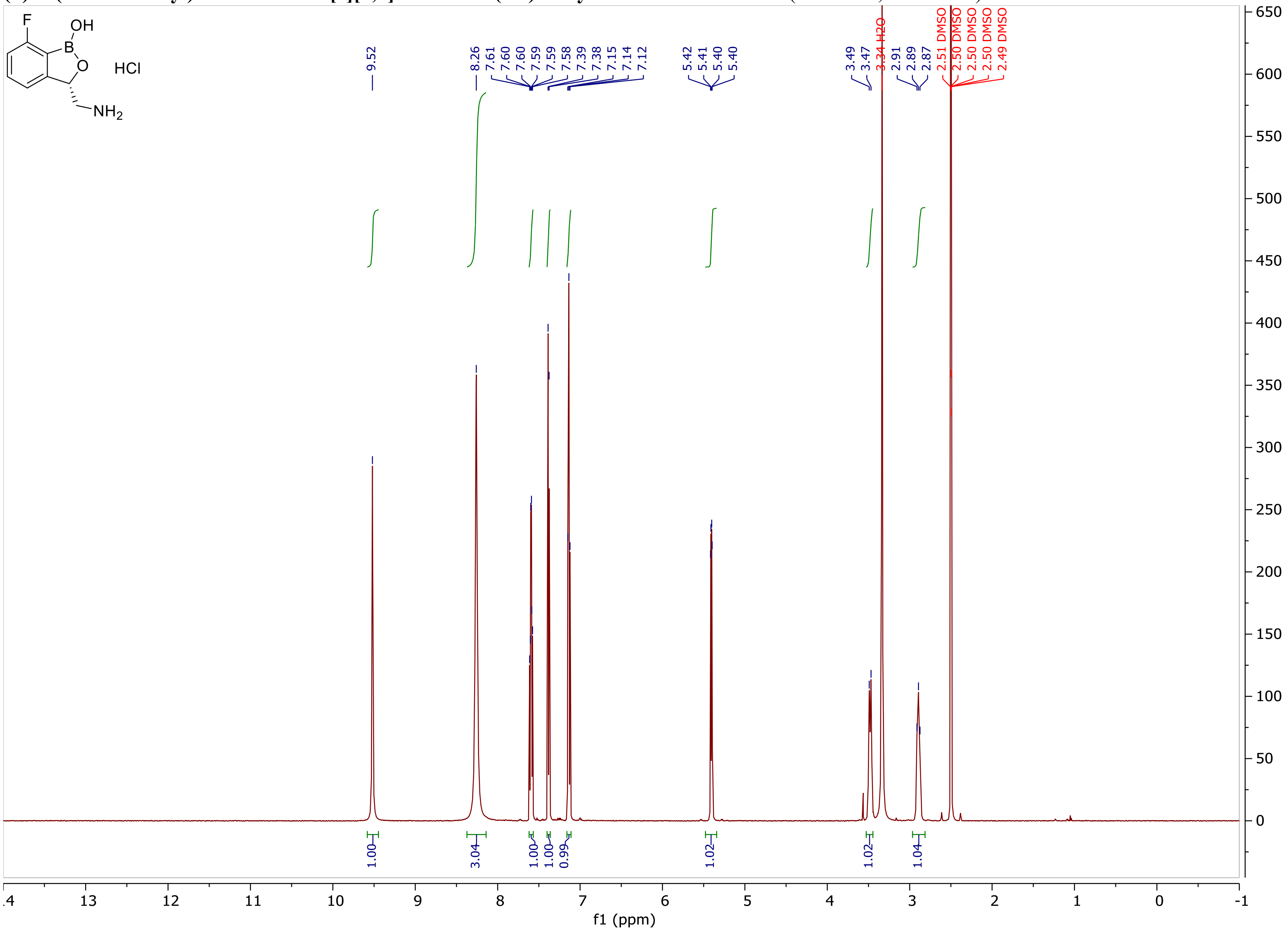

**(*S*)-3-(Aminomethyl)-7-fluorobenzo[*c*][1,2]oxaborol-1(3*H*)-ol hydrochloride.** $^{13}C$ NMR (151 MHz, DMSO-$d_6$)

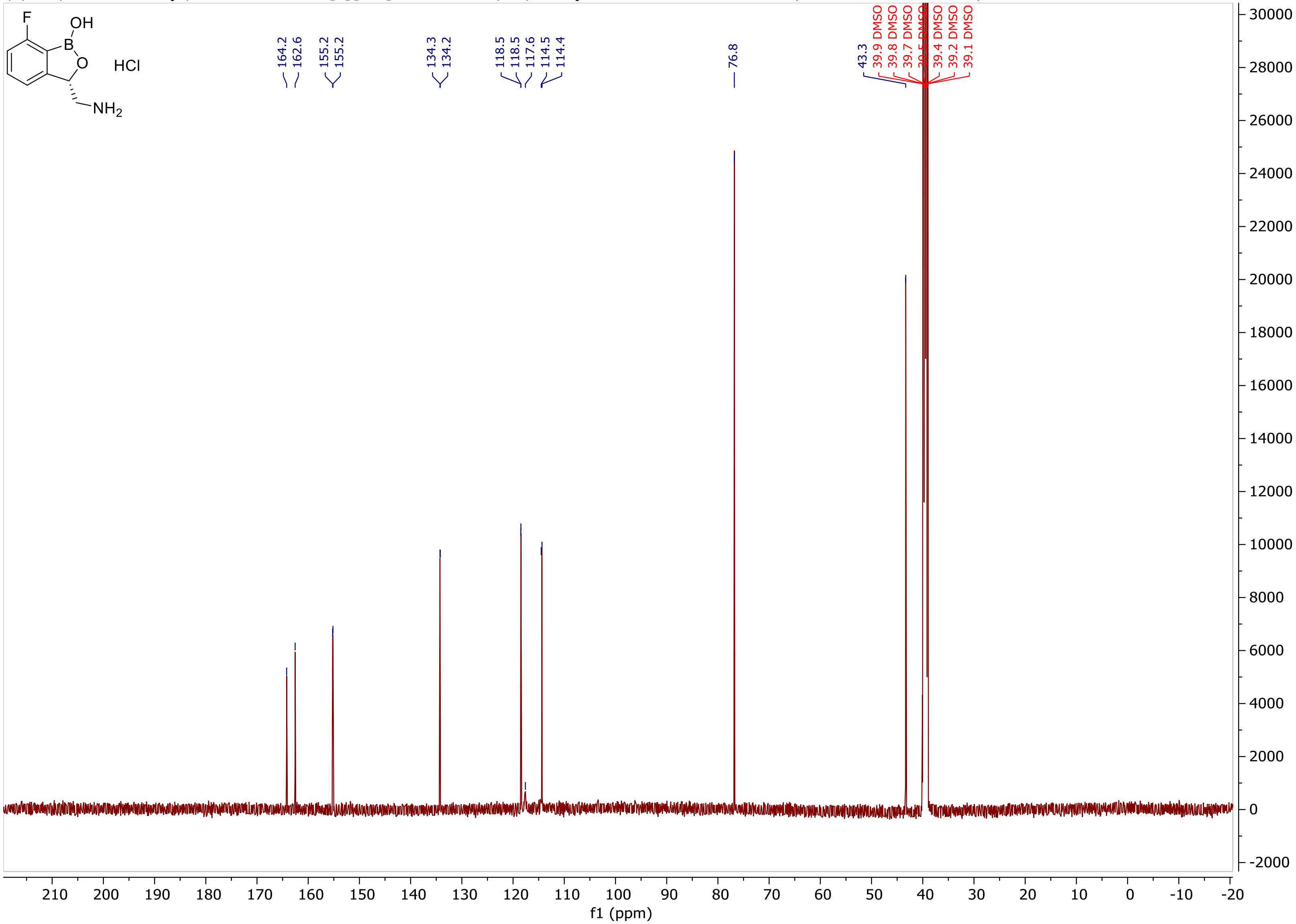

**(*S*)-3-(Aminomethyl)-7-fluorobenzo[*c*][1,2]oxaborol-1(3*H*)-ol hydrochloride.** $^{19}$F NMR (376 MHz, DMSO)

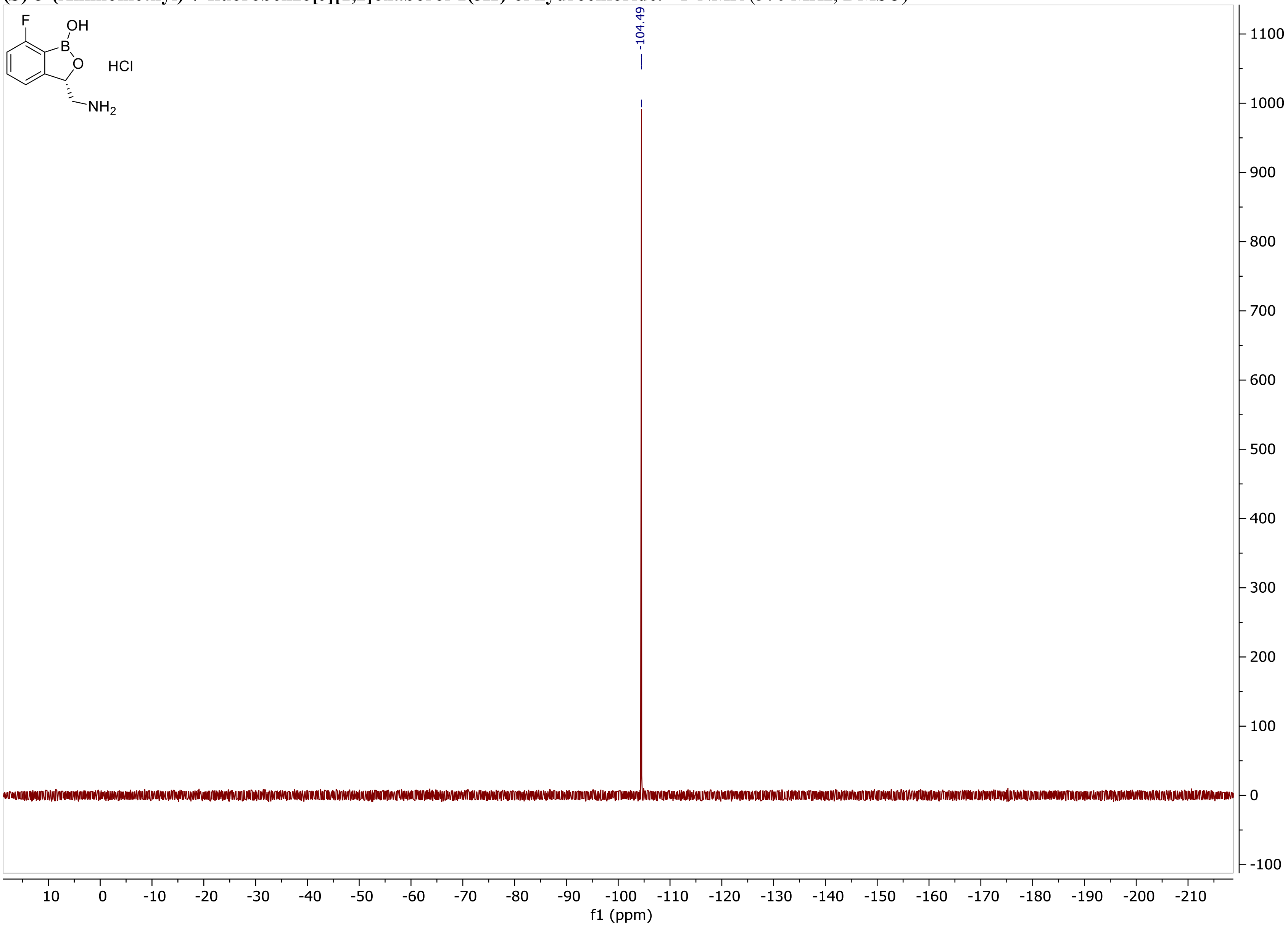

**(*S*)-DMP2.** $^{1}H$ NMR (600 MHz, DMSO-$d_6$)

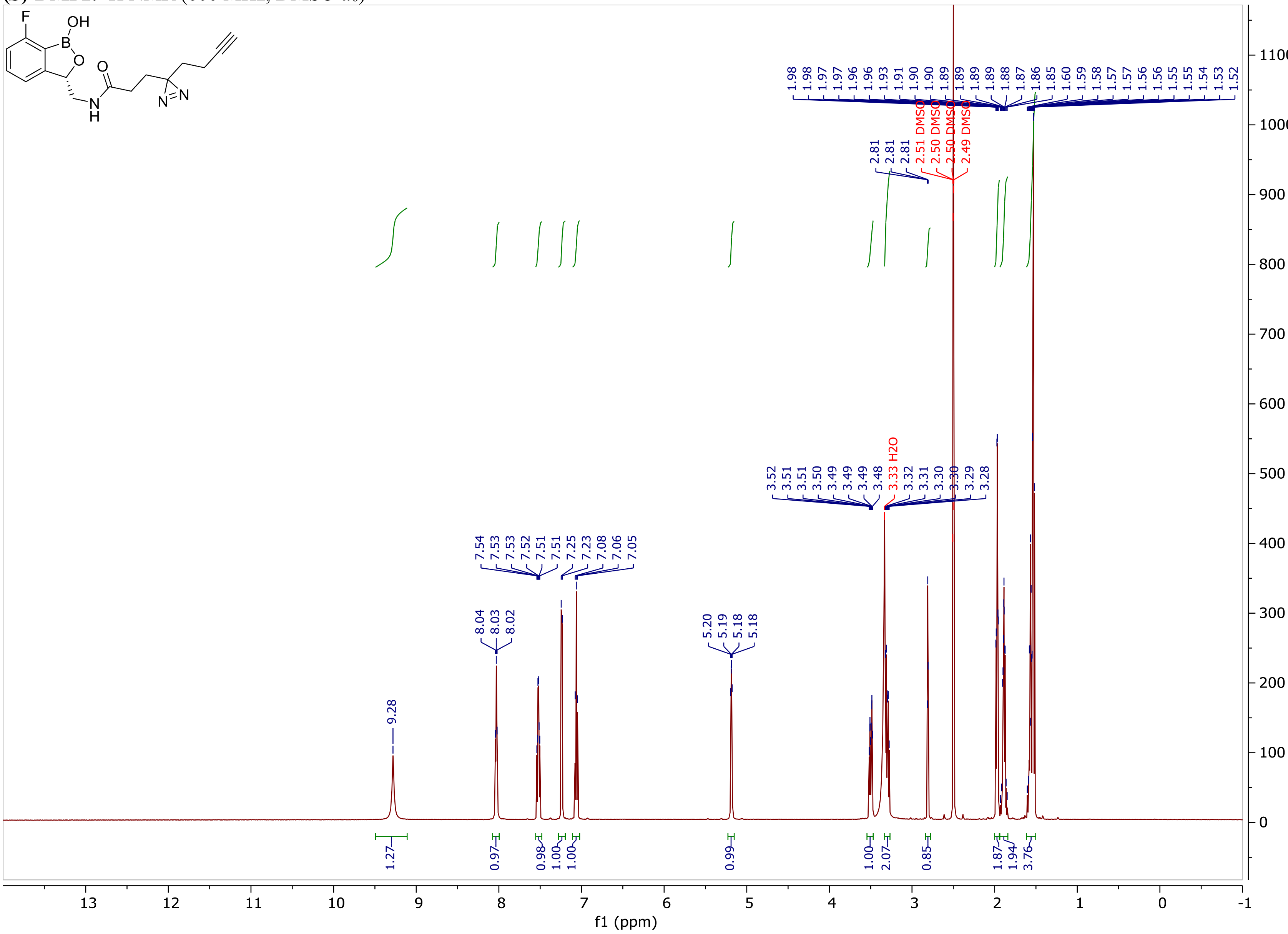

**(*S*)-DMP2.** $^{13}C$ NMR (151 MHz, DMSO-$d_6$)

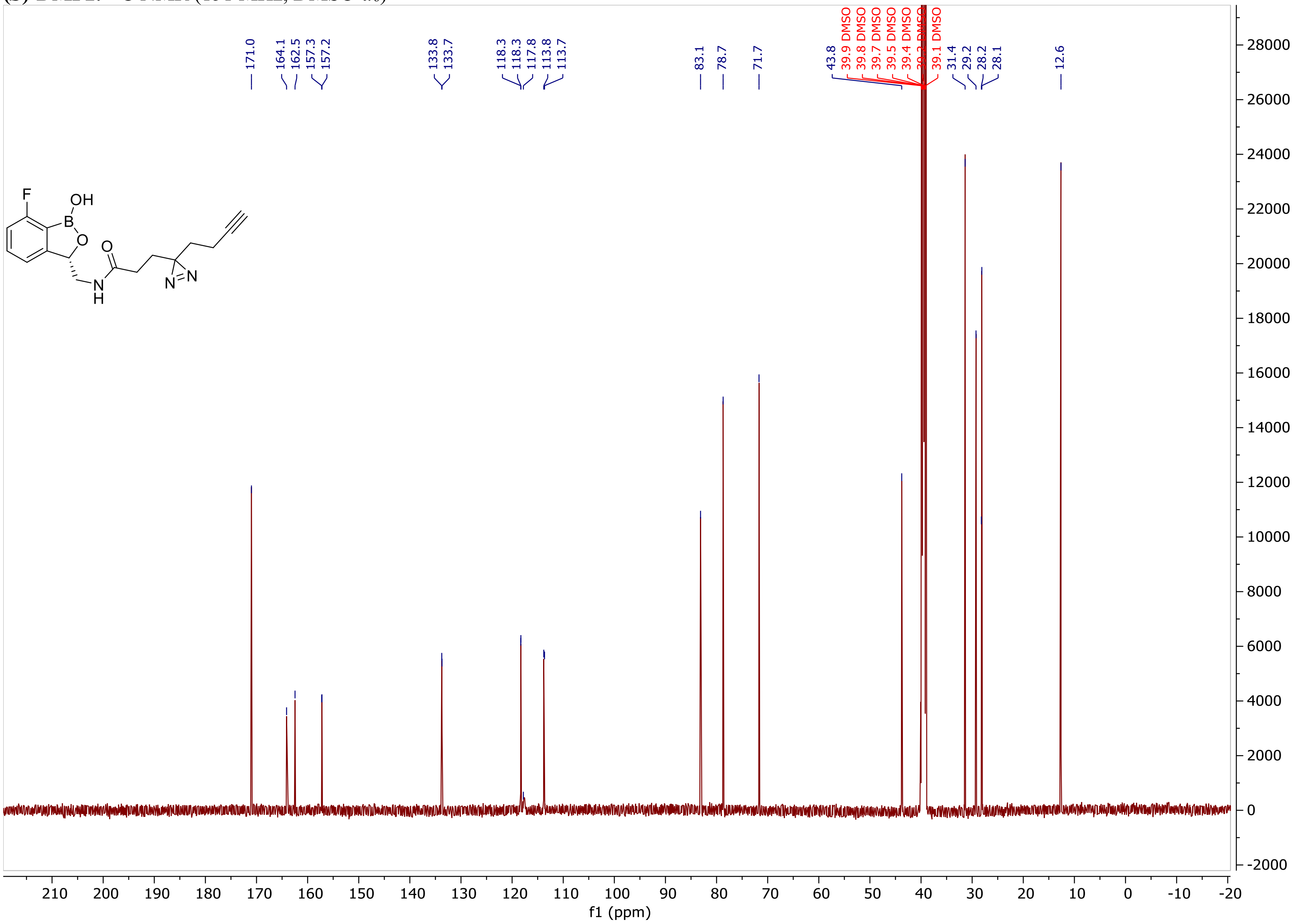

(***S***)**-DMP2.** $^{19}$F NMR (376 MHz, DMSO)

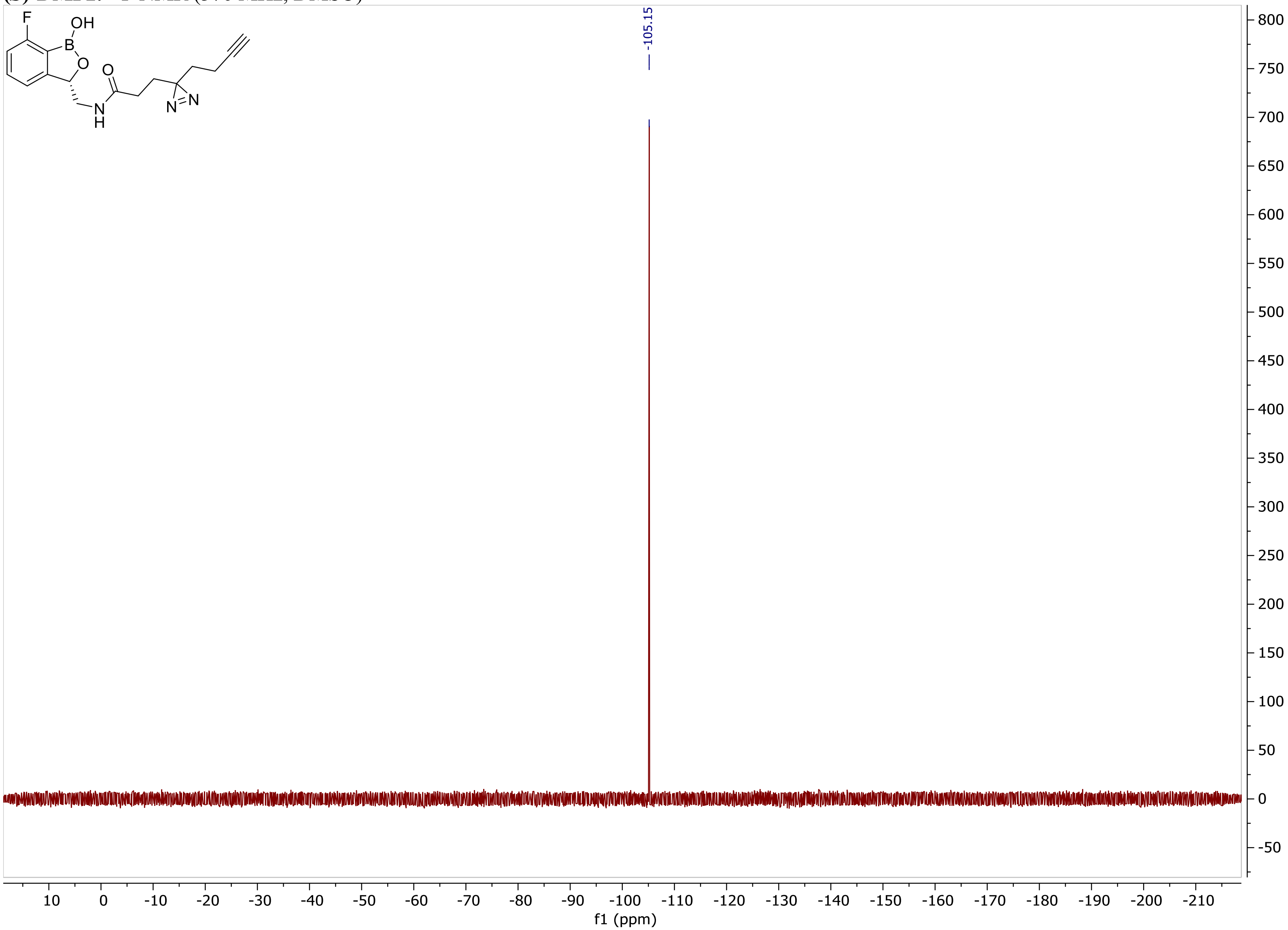

**DMP3.** $^{1}H$ NMR (600 MHz, DMSO-$d_6$)

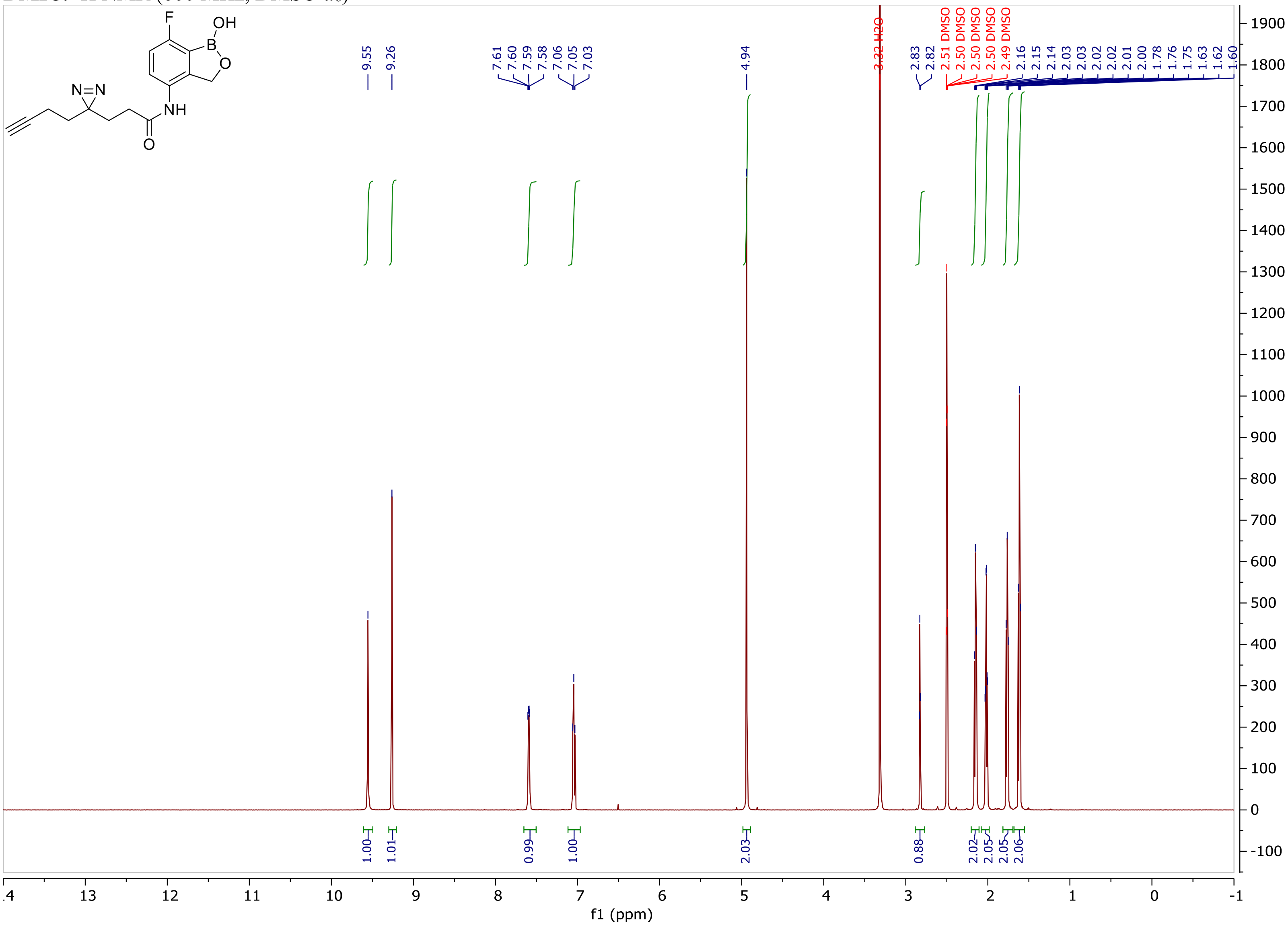

**DMP3.** $^{13}C$ NMR (151 MHz, DMSO-$d_6$)

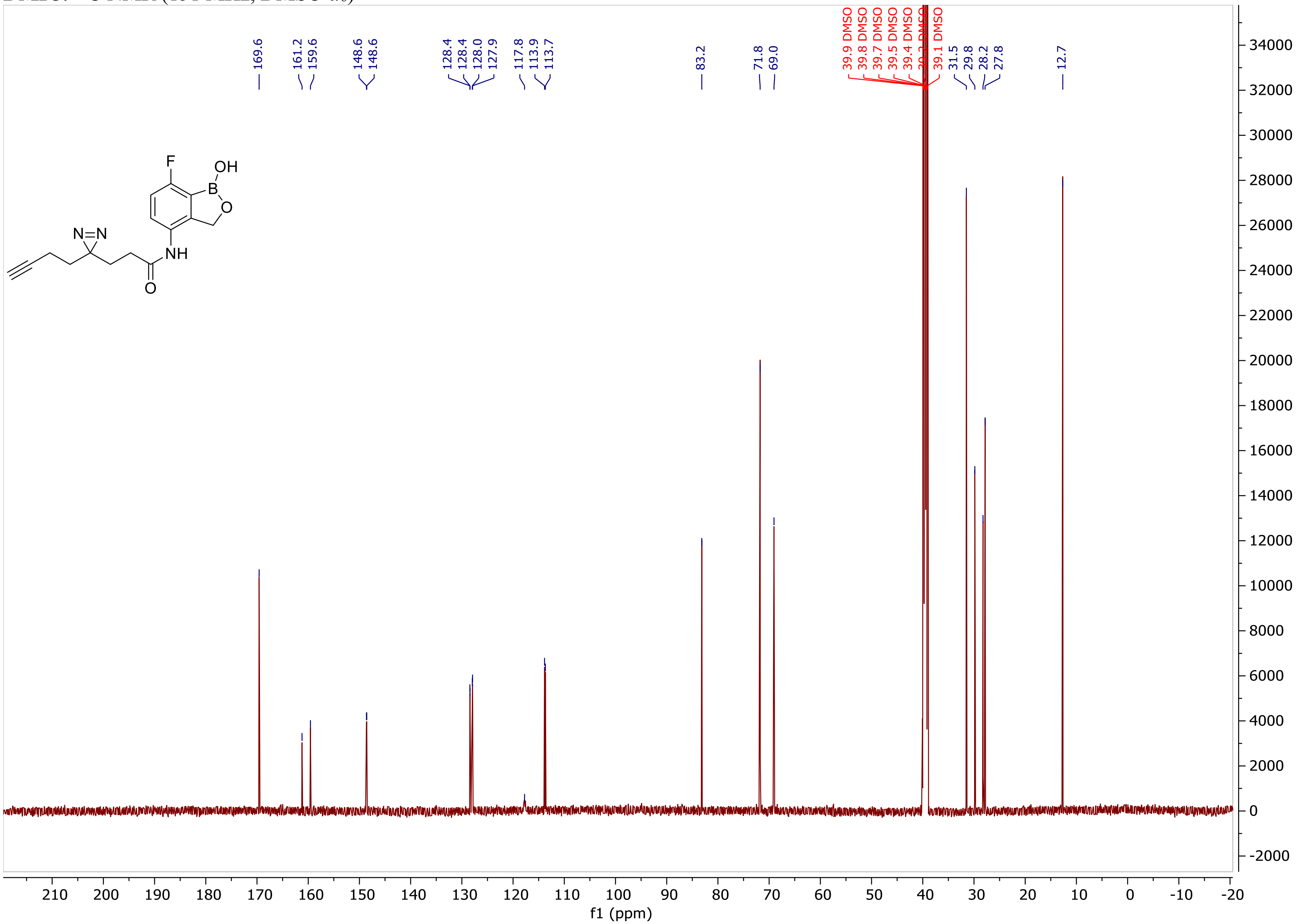

**DMP3.** $^{19}$F NMR (376 MHz, DMSO)

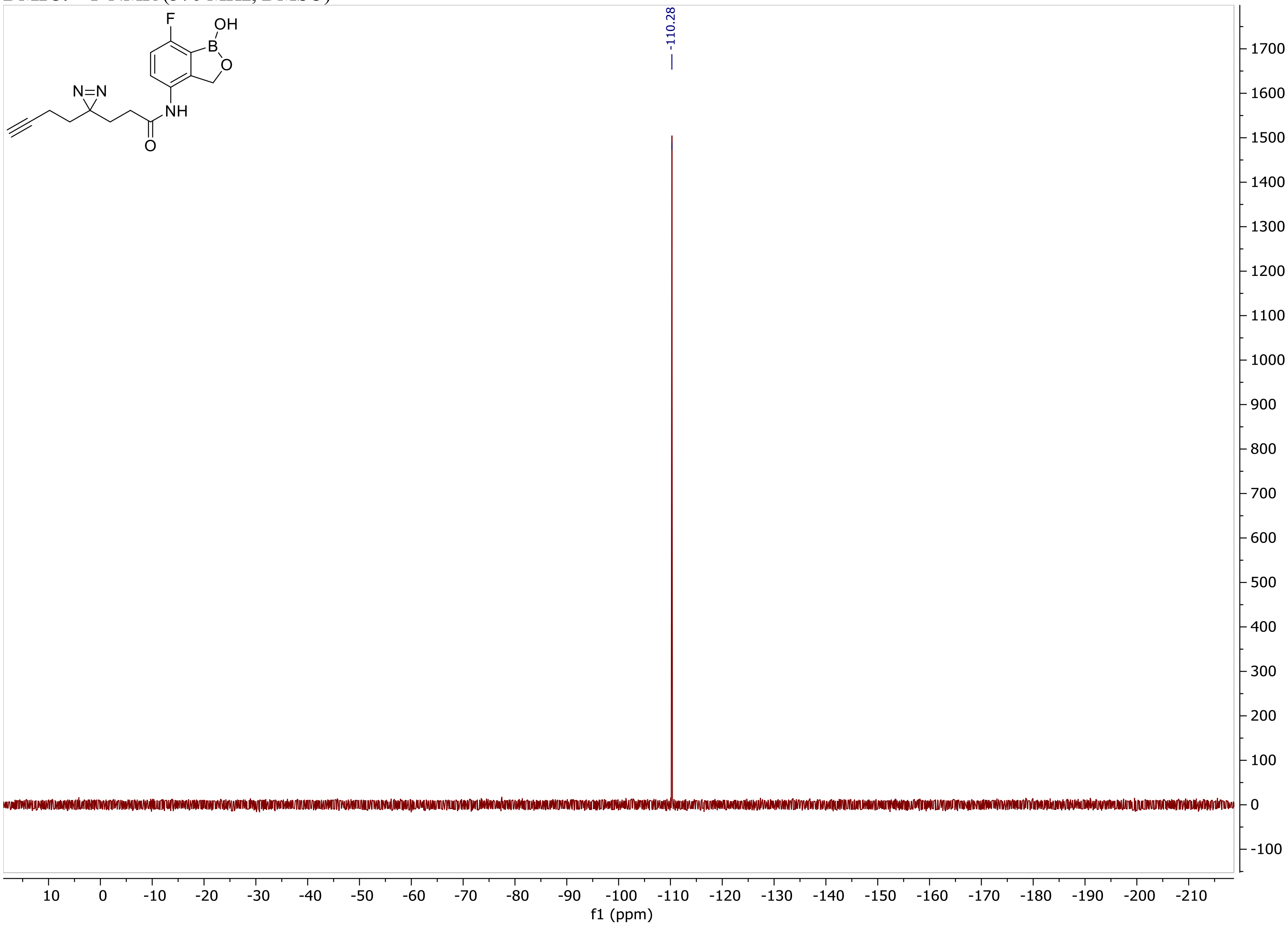

**DMP4.** [1]H NMR (600 MHz, DMSO-$d_6$)

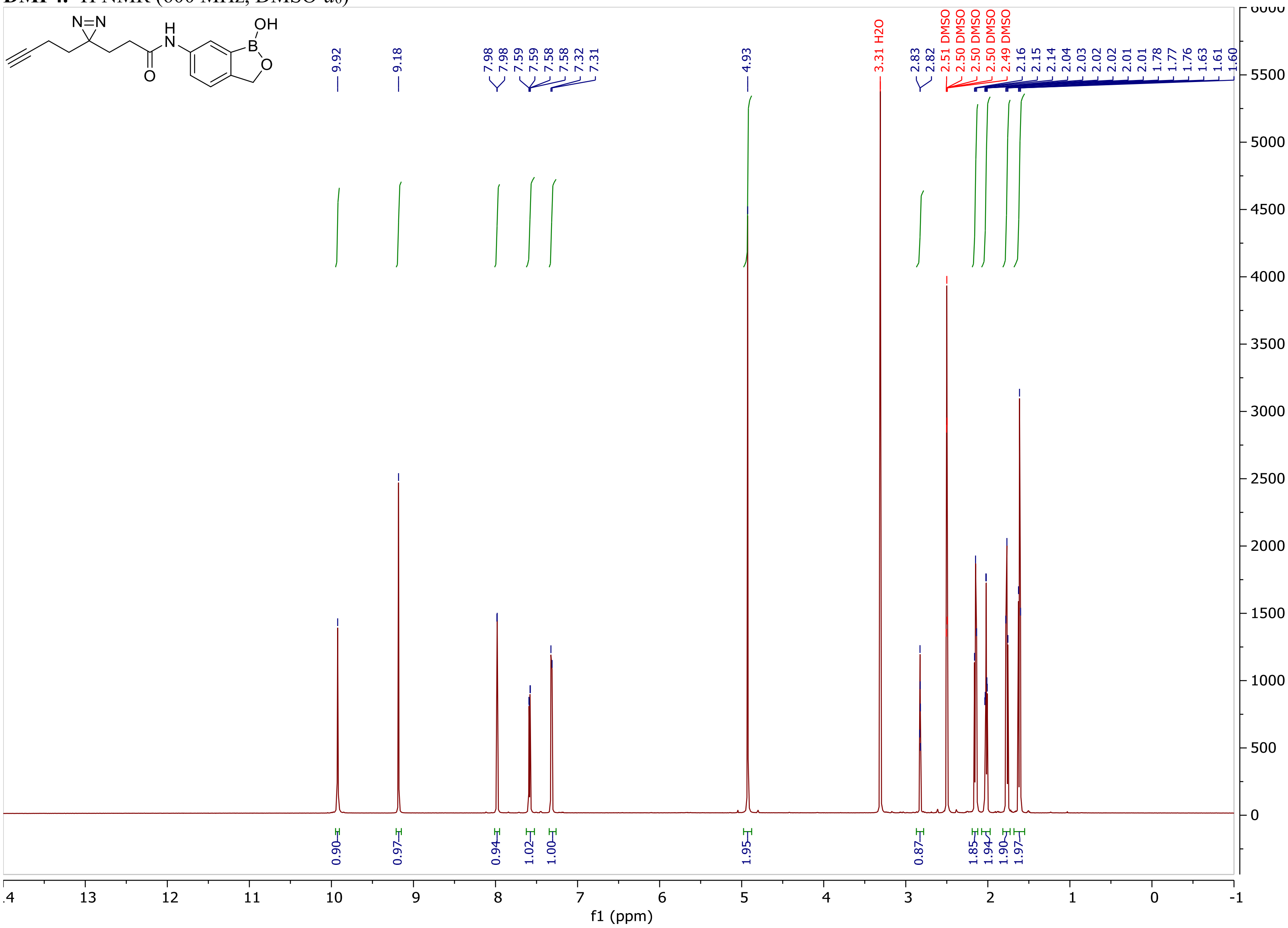

**DMP4.** $^{13}$C NMR (151 MHz, DMSO-$d_6$)

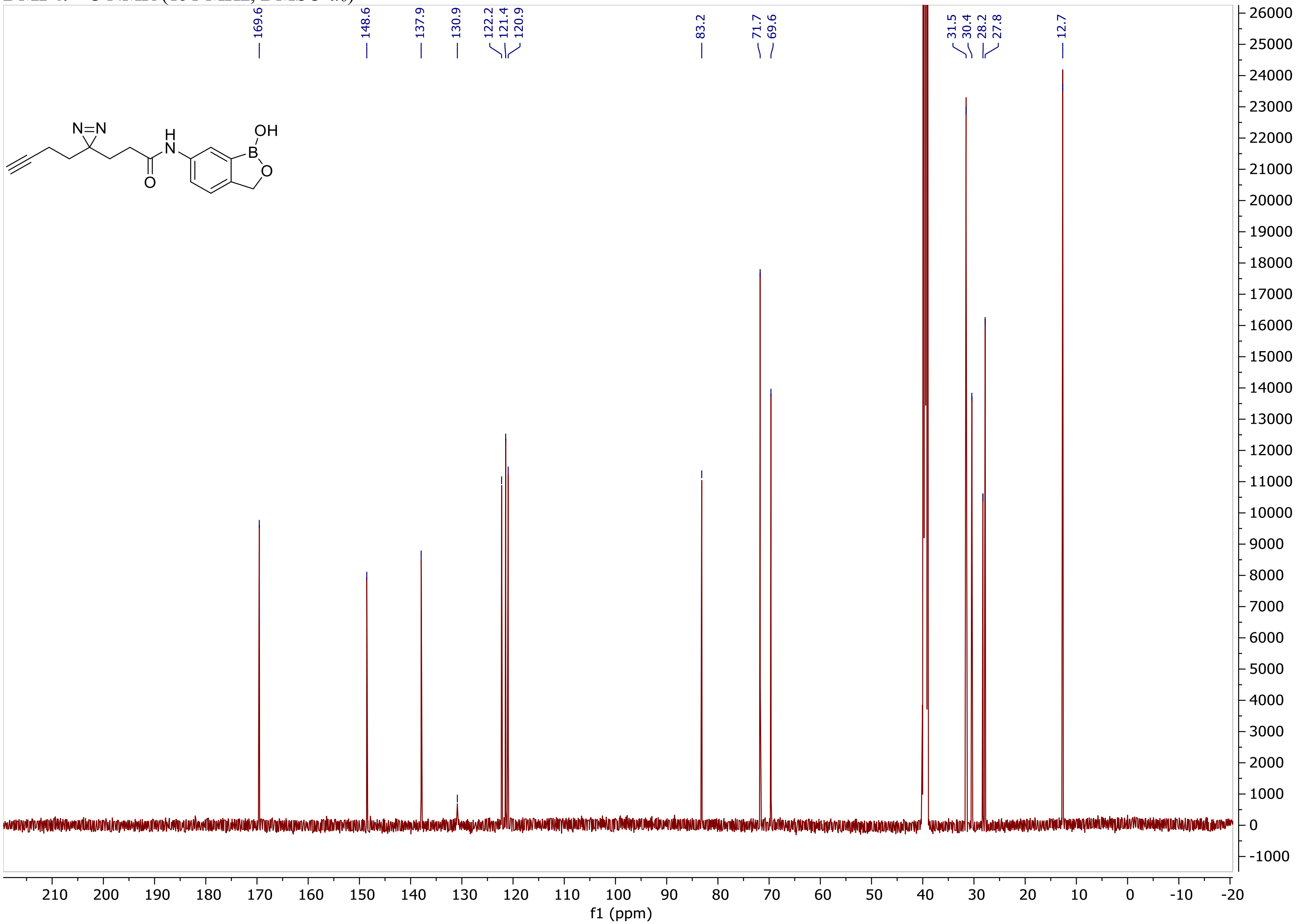

**DMP5.** $^{1}$H NMR (600 MHz, DMSO-$d_6$)

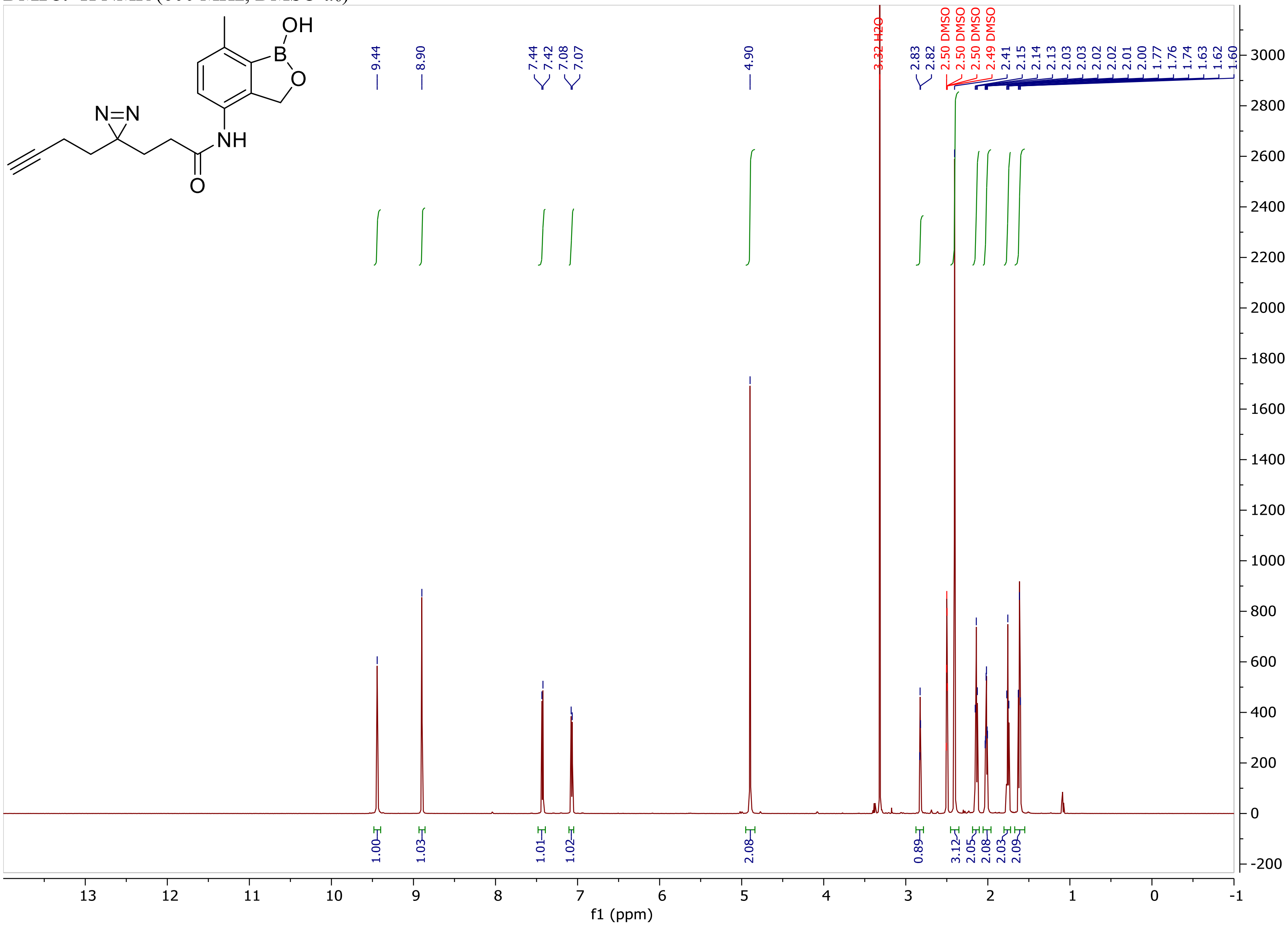

**DMP5.** $^{13}$C NMR (151 MHz, DMSO-$d_6$)

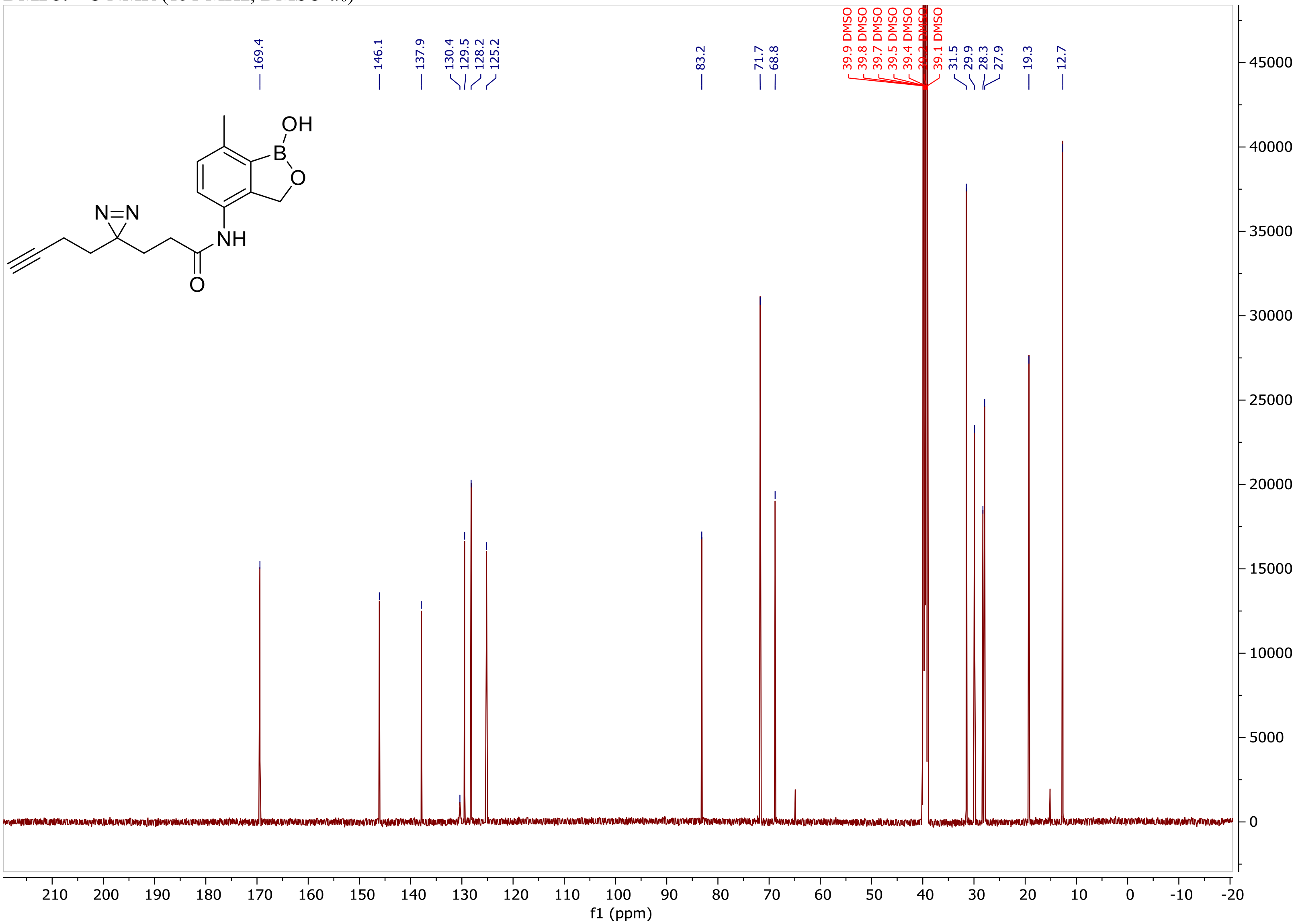

**DMP5.** $^{1}$H COSY NMR (400 MHz, DMSO-$d_6$)

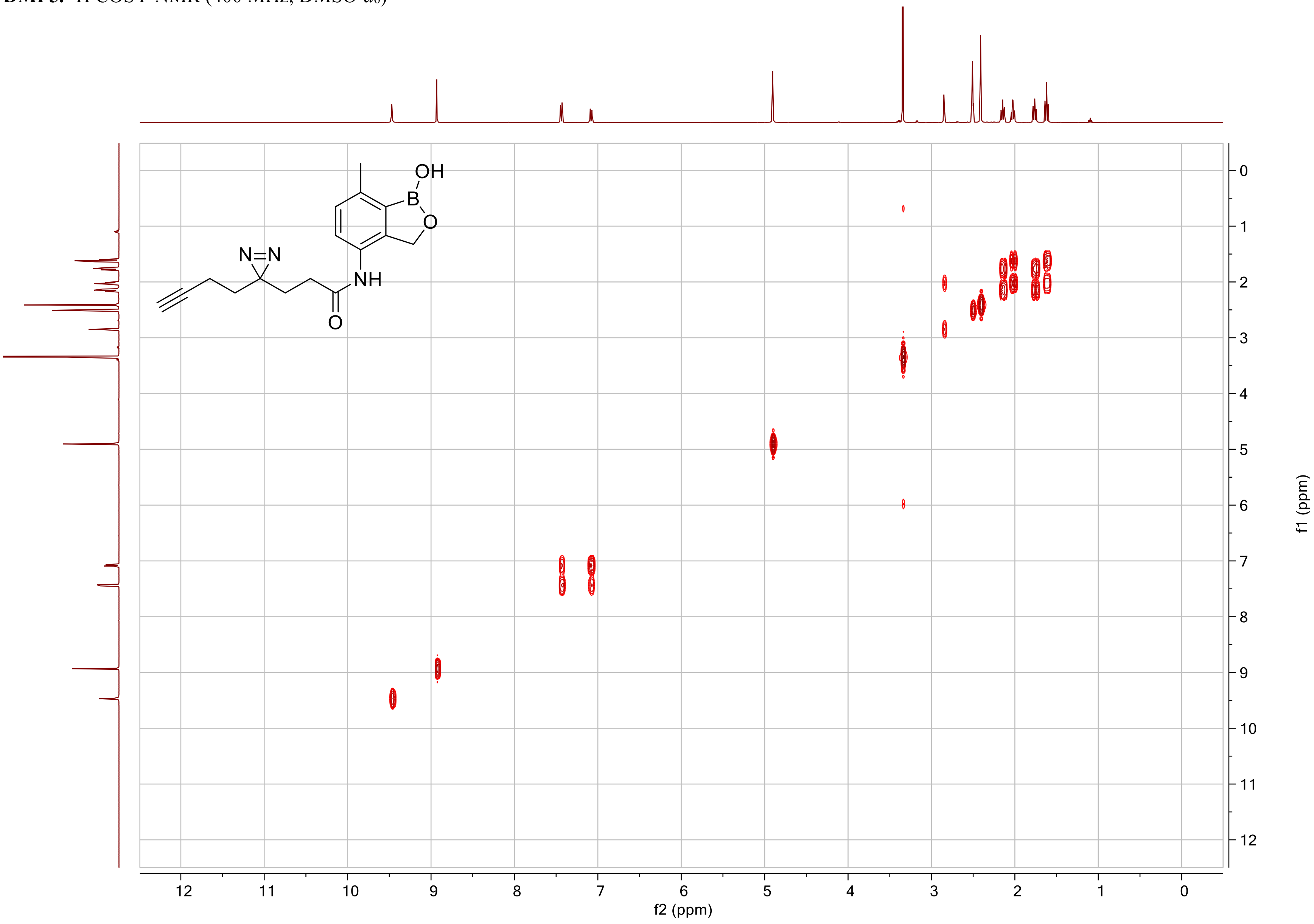

**DMP5.** $^{1}$H HSQC-ed NMR (400 MHz, DMSO-$d_6$)

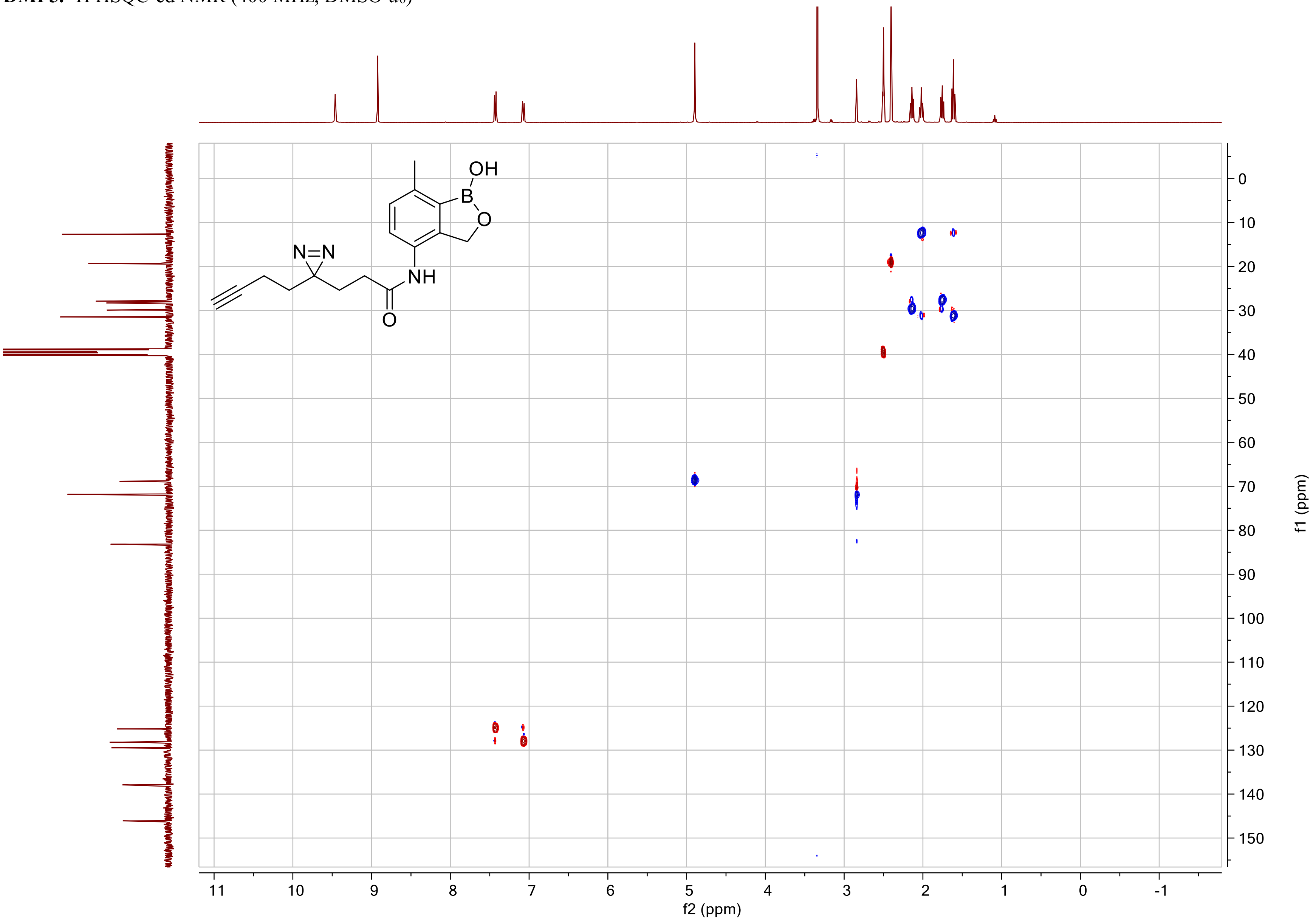

**DMP5.** $^1$H HMBC NMR (400 MHz, DMSO-$d_6$)

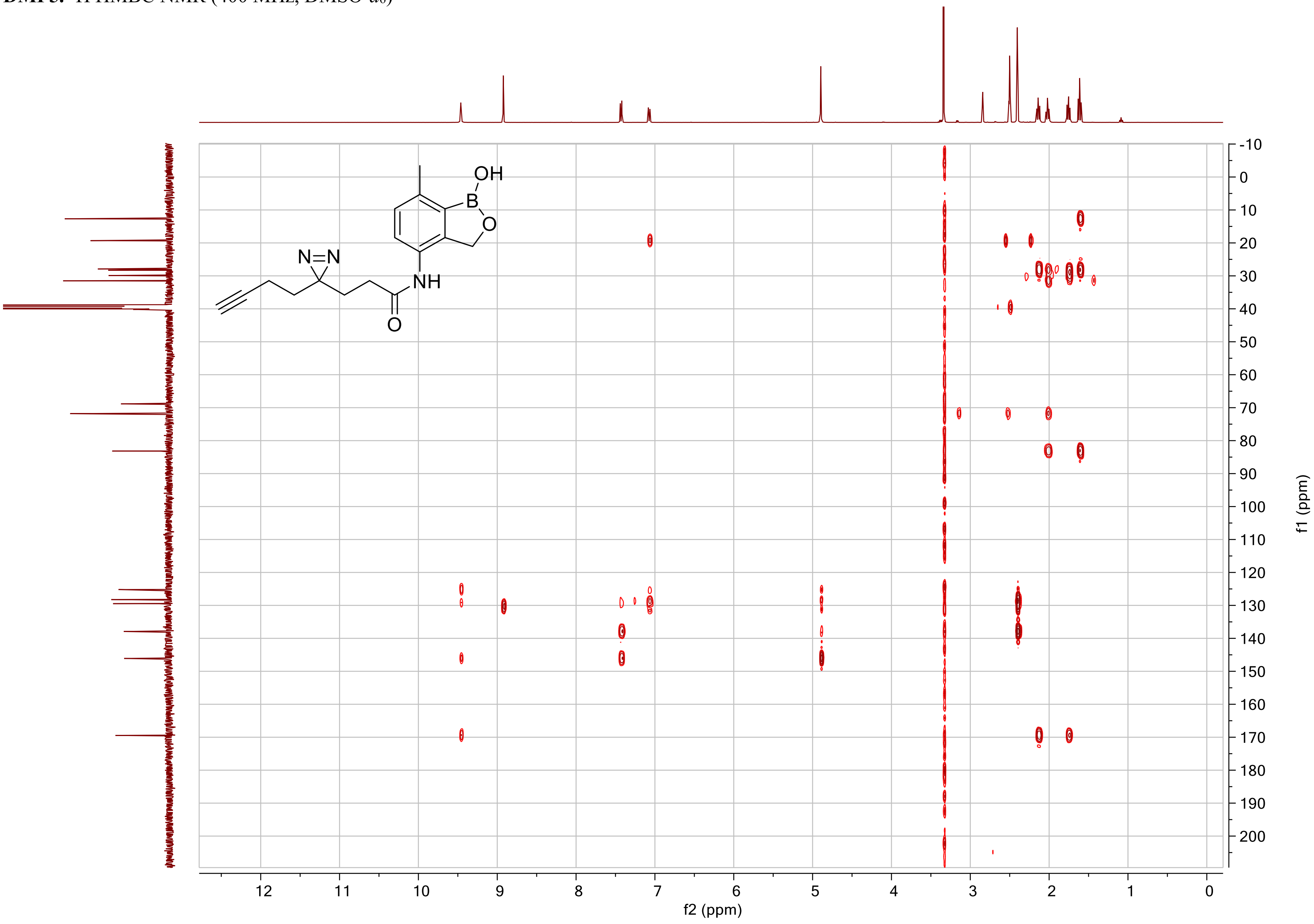

*References*

(1) Wozniak, J. M.; Li, W.; Governa, P.; Chen, L. Y.; Jadhav, A.; Dongre, A.; Forli, S.; Parker, C. G. Enhanced mapping of small-molecule binding sites in cells. *Nat Chem Biol* **2024**, *20* (7), 823-834.
(2) Rock, F. L.; Mao, W.; Yaremchuk, A.; Tukalo, M.; Crepin, T.; Zhou, H.; Zhang, Y. K.; Hernandez, V.; Akama, T.; Baker, S. J.; et al. An antifungal agent inhibits an aminoacyl-tRNA synthetase by trapping tRNA in the editing site. *Science* **2007**, *316* (5832), 1759-1761.
(3) Freund, Y. R.; Akama, T.; Alley, M. R.; Antunes, J.; Dong, C.; Jarnagin, K.; Kimura, R.; Nieman, J. A.; Maples, K. R.; Plattner, J. J.; et al. Boron-based phosphodiesterase inhibitors show novel binding of boron to PDE4 bimetal center. *FEBS Lett* **2012**, *586* (19), 3410-3414.
(4) Akama, T.; Dong, C.; Virtucio, C.; Sullivan, D.; Zhou, Y.; Zhang, Y. K.; Rock, F.; Freund, Y.; Liu, L.; Bu, W.; et al. Linking phenotype to kinase: identification of a novel benzoxaborole hinge-binding motif for kinase inhibition and development of high-potency rho kinase inhibitors. *J Pharmacol Exp Ther* **2013**, *347* (3), 615-625.
(5) Newman, H.; Krajnc, A.; Bellini, D.; Eyermann, C. J.; Boyle, G. A.; Paterson, N. G.; McAuley, K. E.; Lesniak, R.; Gangar, M.; von Delft, F.; et al. High-Throughput Crystallography Reveals Boron-Containing Inhibitors of a Penicillin-Binding Protein with Di- and Tricovalent Binding Modes. *J Med Chem* **2021**, *64* (15), 11379-11394.
(6) Abramson, J.; Adler, J.; Dunger, J.; Evans, R.; Green, T.; Pritzel, A.; Ronneberger, O.; Willmore, L.; Ballard, A. J.; Bambrick, J.; et al. Accurate structure prediction of biomolecular interactions with AlphaFold 3. *Nature* **2024**, *630* (8016), 493-500.
(7) Parker, C. G.; Galmozzi, A.; Wang, Y.; Correia, B. E.; Sasaki, K.; Joslyn, C. M.; Kim, A. S.; Cavallaro, C. L.; Lawrence, R. M.; Johnson, S. R.; et al. Ligand and Target Discovery by Fragment-Based Screening in Human Cells. *Cell* **2017**, *168* (3), 527-541 e529.
(8) Glinka, T.; Higuchi, R.; Hecker, S.; Eastman, B.; Rodny, O. Boron containing polybasic bacterial efflux pump inhibitors and therapeutics uses there of. WO 2012/109164 A1 (2012).
(9) Baker, S. J.; Hernandez, V. S.; Sharma, R.; Nieman, J. A.; Akama, T.; Zhang, Y.-K.; Plattner, J. J.; Alley, M. R. K.; Singh, R.; Rock, F. Boron-containing small molecules. US20090227541A1 (2009).
